# Dynamic regulation of chromatin accessibility by pluripotency transcription factors across the cell cycle

**DOI:** 10.1101/698571

**Authors:** Elias T. Friman, Cédric Deluz, Antonio C.A. Meireles-Filho, Subashika Govindan, Vincent Gardeux, Bart Deplancke, David M. Suter

## Abstract

The pioneer activity of transcription factors allows for opening of inaccessible regulatory elements and has been extensively studied in the context of cellular differentiation and reprogramming. In contrast, the function of pioneer activity in self-renewing cell divisions and across the cell cycle is poorly understood. Here we assessed the interplay between OCT4 and SOX2 in controlling chromatin accessibility of mouse embryonic stem cells. We found that OCT4 and SOX2 operate in a largely independent manner even at co-occupied sites, and that their cooperative binding is mostly mediated indirectly through regulation of chromatin accessibility. Controlled protein degradation strategies revealed that the uninterrupted presence of OCT4 is required for post-mitotic re-establishment and interphase maintenance of chromatin accessibility, and that highly OCT4-bound enhancers are particularly vulnerable to transient loss of OCT4 expression. Our study sheds light on the constant pioneer activity required to maintain the dynamic pluripotency regulatory landscape in an accessible state.

## Introduction

Transcription factors (TFs) regulate the expression of genes through interactions with specific DNA sequences located in gene promoters and distal regulatory elements. A minority of TFs display pioneer activity, i.e. they have the ability to bind and induce the opening of nucleosome-occupied chromatin regions, allowing for the subsequent binding of other TFs and co-factors required for transcriptional activation (Cirillo et al., 2002; Schaffner, 2015; Zaret and Carroll, 2011). Pioneer TFs thereby play a central role in developmental and reprogramming cell fate decisions, which hinge on large-scale reshaping of the chromatin landscape in tissue-specific regulatory regions (Chronis et al., 2017; Iwafuchi-Doi and Zaret, 2014; Jacobs et al., 2018; Mayran et al., 2018; Pastor et al., 2018; Soufi et al., 2015, 2012; Takaku et al., 2016). Much less is known about the role of pioneer activity and of its dynamics over the cell cycle in regulating stem cell self-renewal. The OCT4 (also known as POU5F1) and SOX2 pioneer TFs (Soufi et al., 2015) are absolutely required for the self-renewal of embryonic stem (ES) cells (Masui et al., 2007; Niwa et al., 2000). OCT4 and SOX2 can form a heterodimer that binds to a composite motif at thousands of sites in the genome (Boyer et al., 2005; Nishimoto et al., 1999; Yuan et al., 1995). A recent study has shown that depletion of OCT4 for 24 hours in ES cells leads to loss of accessibility and co-factor occupancy at a large fraction of its bound enhancers involved in pluripotency maintenance (King and Klose, 2017). In contrast, the role of SOX2 in the regulation of ES cell chromatin accessibility has not been elucidated. Thus, to which extent the pioneering activities of OCT4 and SOX2 overlap and/or depend on each other to regulate chromatin accessibility in ES cells is unclear.

Self-renewal requires the ability to progress through the cell cycle without losing cell type-specific gene expression. This is not a trivial task since chromatin accessibility of gene regulatory elements is markedly decreased during S phase and mitosis (Festuccia et al., 2019; Hsiung et al., 2015; Oomen et al., 2019; Stewart-Morgan et al., 2019). How recovery of chromatin accessibility after DNA replication and mitosis is controlled, and whether it requires pioneer activity is poorly understood. The period of genome reactivation occurring at the mitosis-G1 (M-G1) transition coincides with a particularly favorable context for reprogramming by somatic cell nuclear transfer (mitosis) (Egli et al., 2008) and increased sensitivity to differentiation signals in human ES cells (G1 phase) (Pauklin and Vallier, 2013). Recent evidence also points at cell cycle stage-specific functions of OCT4 and SOX2 in cell fate regulation. OCT4 expression levels in G1 phase affect the propensity of ES cells to differentiate towards neuroectoderm and mesendoderm (Strebinger et al., 2019), and depletion of OCT4 at the M-G1 transition impairs pluripotency maintenance of ES cells and leads to a lower reprogramming efficiency upon overexpression in mouse embryonic fibroblasts (Liu et al., 2017). Depletion of SOX2 at the M-G1 transition impairs both pluripotency maintenance and SOX2-induced neuroectodermal differentiation of ES cells upon release of pluripotency signals (Deluz et al., 2016). Whether the particular sensitivity of M and G1 phases to the action of OCT4 and SOX2 is related to the dynamics of their pioneer activity across the cell cycle is unknown.

Here we studied the interplay of OCT4 and SOX2 in regulating chromatin accessibility of ES cells and dissected the pioneer activity of OCT4 across the cell cycle. We show that most enhancers bound by both TFs depend on only one of them to maintain their open chromatin state, and that cooperative binding of OCT4 and SOX2 is mainly mediated indirectly through changes in chromatin accessibility. Using forms of OCT4 engineered for mitotic or auxin-inducible degradation, we demonstrate the role of OCT4 in re-establishment and continuous maintenance of chromatin accessibility throughout the cell cycle.

## Results

### OCT4 and SOX2 regulate chromatin accessibility at mostly distinct loci

OCT4 and SOX2 bind cooperatively to thousands of genomic locations in ES cells both independently and as a heterodimer on a composite OCT4::SOX2 motif. How OCT4 and SOX2 interplay to regulate chromatin accessibility in ES cells is not known. To address this question, we decided to determine genome-wide chromatin accessibility changes upon acute loss of OCT4 or SOX2. To deplete OCT4 and SOX2 from ES cells in an inducible manner, we took advantage of the ZHBTc4 (Niwa et al., 2000) and 2TS22C (Masui et al., 2007) mouse ES cell lines, in which a Tet-Off promoter regulates the expression of *Oct4* and *Sox2*, respectively (Fig.1A). While OCT4 is fully depleted after 24 hours of doxycycline (dox) (Niwa et al., 2000), SOX2 is not, likely due to its longer half-life (Masui et al., 2007). We determined SOX2 levels by immunofluorescence staining after 26 and 40 hours of dox treatment and found that residual SOX2 expression persisted after 26 hours but not 40 hours of dox treatment (Fig. S1A). Importantly, despite its known role in regulating expression of OCT4 (Dunn et al., 2014; Strebinger et al., 2019), SOX2 depletion for 26 or 40 hours had only a minor impact on OCT4 levels (Fig. S1A-B). In ZHBTc4 cells, as expected 24 hours of dox treatment led to the complete depletion of OCT4 and only mildly affected SOX2 levels (Fig. S1C-D). Therefore, changes in chromatin accessibility upon short-term SOX2 or OCT4 loss are unlikely to be confounded by changes in expression levels of OCT4 and SOX2, respectively.

**Figure 1.**
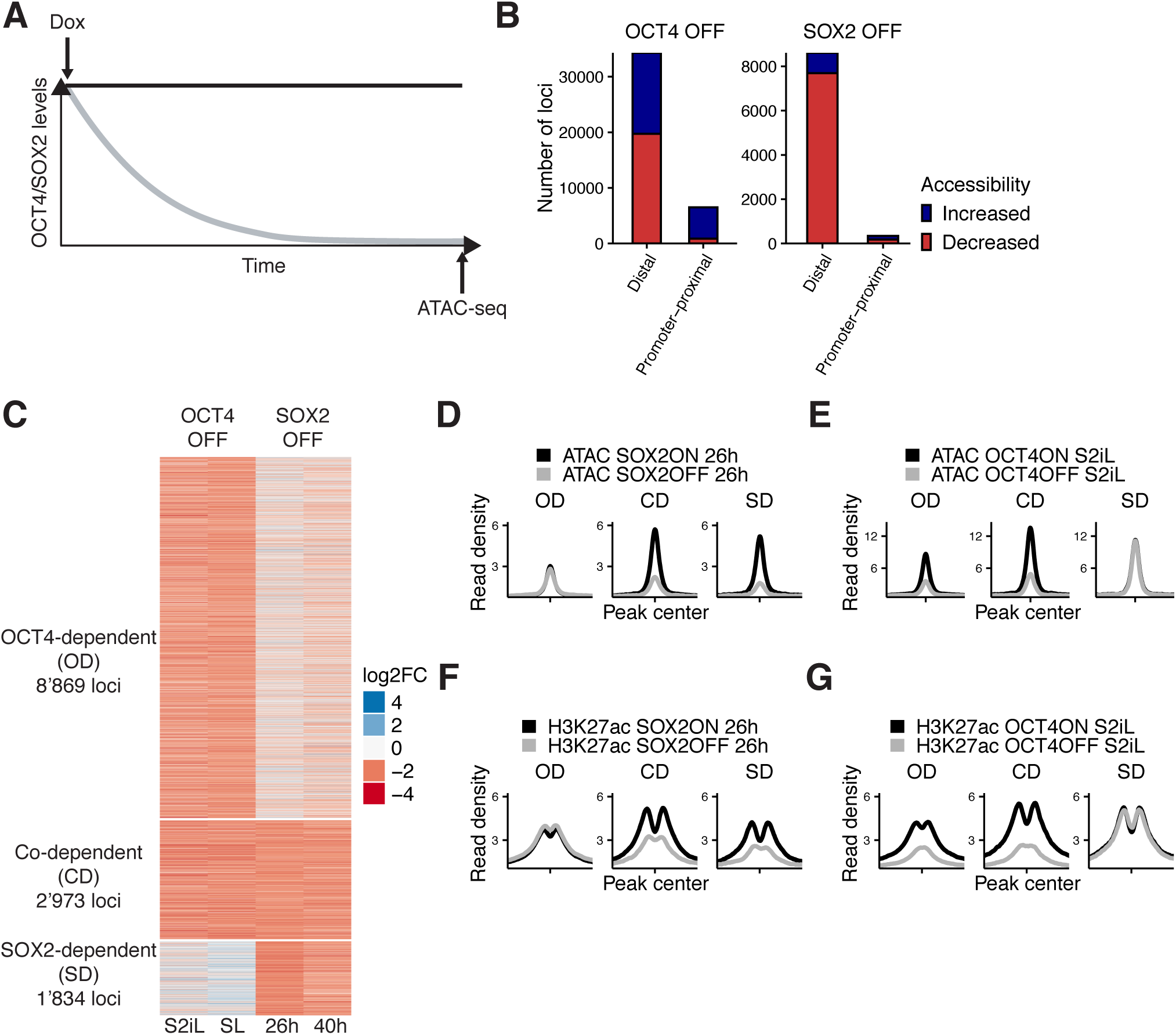
Interplay between OCT4 and SOX2 in regulating ES cell chromatin accessibility. **(A)** Experimental strategy to compare the effect of OCT4 and SOX2 depletion on chromatin accessibility. (B) Number of regions significantly changed in accessibility upon OCT4 (left) and SOX2 (right) depletion in distal (>1 kb from TSS) and promoter-proximal (<1 kb from TSS) elements. (C) log2 fold-change values of accessibility between dox-treated and untreated cells upon OCT4/SOX2 depletion at OCT4/SOX2 binding sites with decreased accessibility. Loci are grouped into those significantly affected upon OCT4 depletion (OD), SOX2 depletion (SD), or depletion of either factor (CD). Each row corresponds to one individual locus, and each column to a different experimental condition. (D-E) Average RPKM-normalized ATAC-seq signal 2 kb around OD, CD, and SD loci upon SOX2 depletion (D) and OCT4 depletion (E). (F-G) Average RPKM-normalized H3K27ac ChIP-seq signal 2 kb around OD, CD, and SD loci upon SOX2 depletion (F) and OCT4 depletion (G).

We performed ATAC-seq in ZHBTc4 cells without dox or with dox for 24 hours, and in 2TS22C cells without dox or with dox for 26 or 40 hours. We first compared chromatin accessibility changes between ZHBTc4 cells +/-dox for 24 hours in our culture conditions (serum + LIF + 2i (S2iL), see Methods) to a previous dataset acquired with ZHBTc4 cells +/-dox for 24 hours but cultured in serum + LIF (SL) (King and Klose, 2017). The good correlation (Pearson’s R=0.7) in chromatin accessibility changes at OCT4 binding sites between culture conditions (Fig. S1E) prompted us to take advantage of both datasets for further analysis. We next compared changes in accessibility at SOX2 binding sites in the 2TS22C cell line treated for either 26 or 40 hours with dox, which also displayed a clear correlation (Pearson’s R=0.61) (Fig.S1F). We reasoned that the 26 hour dox dataset should be less prone to changes in accessibility due to indirect effects of prolonged SOX2 depletion than the 40 hour dox dataset, while the latter should be more sensitive to identify loci that are still accessible at low SOX2 concentrations. We thus called significantly affected loci using limma (Ritchie et al., 2015) (false discovery rate (FDR) < 0.05) and selected only those in which the direction of change (decrease or increase in accessibility) was the same for 26 hours vs 40 hours of dox treatment in 2TS22C cells, and likewise for SL vs S2iL in ZHBTc4 cells. In line with previous reports, loss of OCT4 led to decreased accessibility at 20’587 loci, most of which are distal regulatory elements (Fig. 1B). Loss of SOX2 also led to decreased accessibility mainly at distal elements, but at fewer loci (7’874). We also found that loss of OCT4 led to a gain in accessibility at 20’209 loci, while 1’080 loci gained accessibility upon loss of SOX2 (Fig. 1B). Loci that lost accessibility were highly enriched for OCT4 and SOX2 ChIP-seq binding while loci that gained accessibility were much less so (Fig.S2A-B).

To compare the loci impacted by OCT4 vs SOX2 depletion, we next focused on all regions that were bound by OCT4 and/or SOX2 as identified from available and newly generated ChIP-seq datasets (see Fig. S2A-B and Methods) and that lost accessibility upon dox treatment. To avoid misrepresenting differences in SOX2 and OCT4 regulation that arise from differences in accessibility due to culture conditions or cell lines, we called significantly different loci (FDR < 0.05) between untreated ZHBTc4 cells cultured in SL vs S2iL conditions as well as between untreated ZHBTc4 cells and 2TS22C cells in S2iL. We then discarded all loci that displayed a large difference (FC > 4) in any of those comparisons. We classified the remaining loci as OCT4-dependent (OD, 8’869 loci), SOX2-dependent (SD, 1’834 loci), and co-dependent (CD, 2’973 loci), as defined by loss of accessibility upon depletion of OCT4 only, SOX2 only, or either of them, respectively (Fig. S3A, Fig. 1C-E). All three groups were enriched for chromatin marks of enhancers (Fig. S3B). We performed ChIP-seq analysis of the active enhancer mark H3K27ac (Creyghton et al., 2010) upon OCT4 or SOX2 loss for 24 hours and 26 hours, respectively. All groups displayed a reduction in H3K27ac, suggesting concordant maintenance of enhancer accessibility and activity by OCT4 and/or SOX2 at these loci (Fig.1F-G).

**Figure 3.**
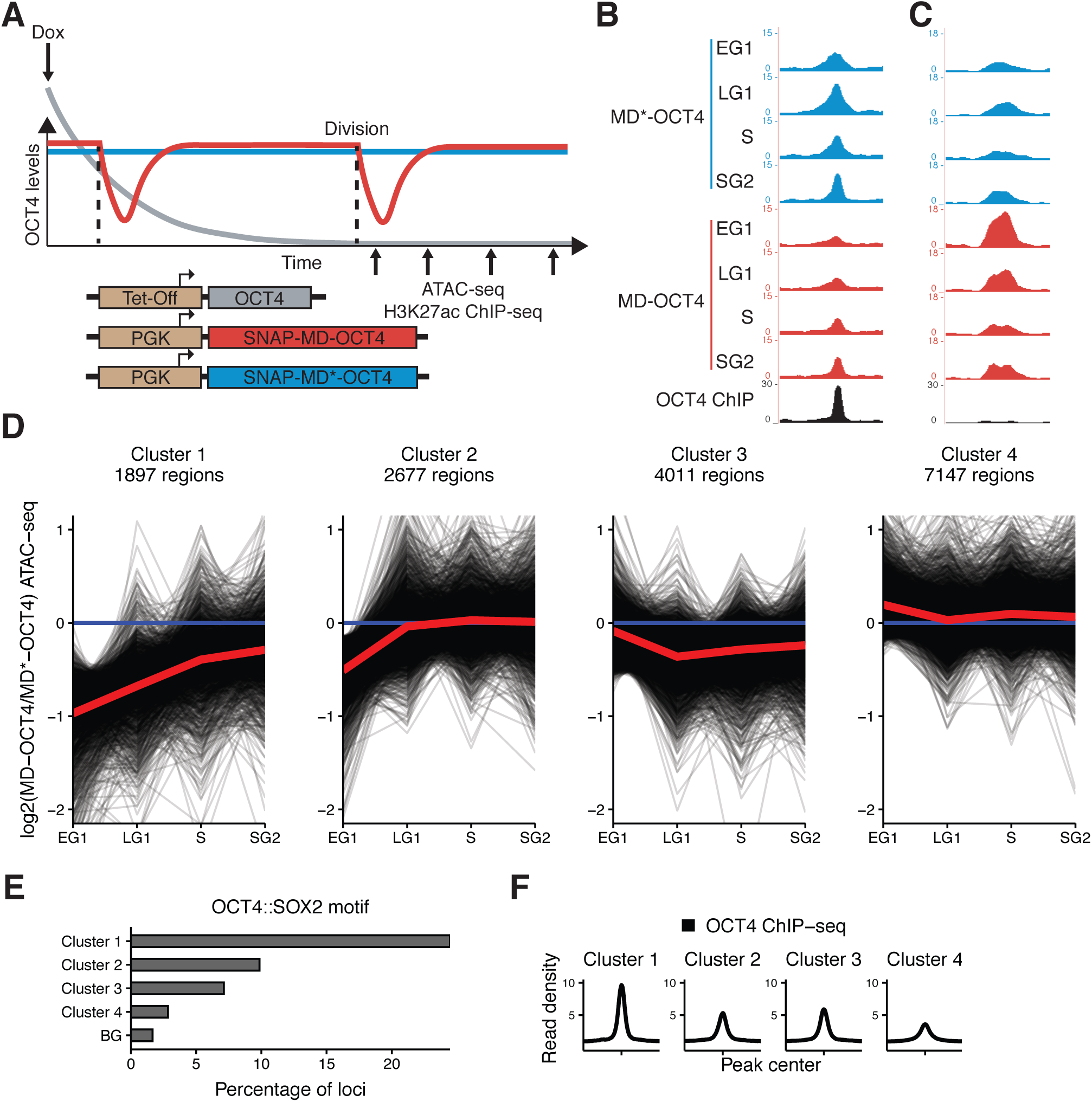
Mitotic degradation of OCT4 results in different patterns of accessibility loss. **(A)** Experimental strategy used to assess the impact of OCT4 depletion at the M-G1 transition. (B-C) Genome browser tracks of RPKM-normalized accessibility profiles across the cell cycle for one locus that decreases (B) at chr11:6894809-6895533 and one that increases (C) at chr9:41247953-4124841 in accessibility upon transient OCT4 depletion in M-G1. (D) log2 fold-change values of accessibility between MD-OCT4 and MD*-OCT4 (control) cells in different cell cycle phases at all accessible OCT4-bound sites, grouped into four clusters by k-means clustering (see Methods). Each line represents one locus. Red line: mean of each cluster. (E) Frequency of overlap with the canonical OCT4::SOX2 motif in the four clusters as well as in background regions (BG). (F) Average RPKM-normalized OCT4 ChIP-seq signal in untreated ZHBTc4 cells 2 kb around loci in the four clusters.

Surprisingly, all groups were enriched for the binding of both OCT4 and SOX2 (Fig. 2A). 89% of SD sites overlapped with an OCT4 peak and 65% of OD sites overlapped with a SOX2 peak. Therefore, differences in the regulation of chromatin accessibility at these loci cannot simply be explained by differential DNA binding of SOX2 and OCT4. OCT4 has been shown to regulate chromatin accessibility by recruitment of the BAF chromatin remodeling complex, including the BRG1 subunit (King and Klose, 2017). As SOX2 also interacts with BRG1 in vivo (Xu et al., 2018), we asked whether SOX2 also regulates chromatin accessibility through BRG1 recruitment. We performed BRG1 ChIP-seq upon SOX2 depletion and reanalyzed ChIP-seq data of BRG1 upon OCT4 depletion (King and Klose, 2017). We found that loss of accessibility was accompanied by loss of BRG1 in all groups (Fig. 2B-C). We also reanalyzed ATAC-seq data from cells in which BRG1 has been depleted (Ho et al., 2011; King and Klose, 2017) and found that all groups were dependent on BRG1 to maintain their accessibility (Fig. S4A). This suggests that OCT4 and SOX2 can regulate chromatin accessibility independently of each other even at sites that are co-occupied and through the recruitment of BRG1.

**Figure 2.**
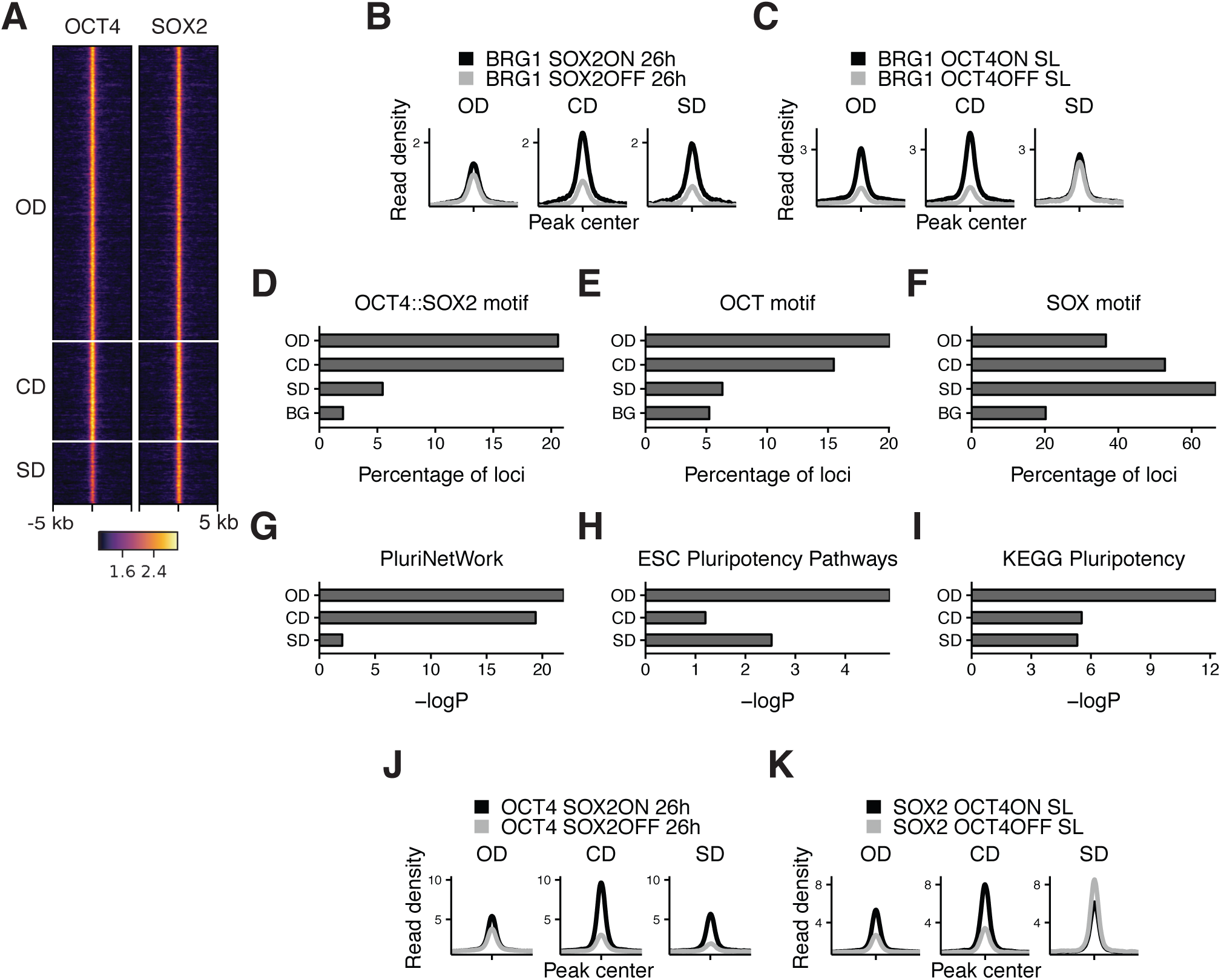
Characterization of OCT4/SOX2-dependent loci. (A) Heatmap of RPKM-normalized OCT4 and SOX2 ChIP-seq binding profiles in untreated ZHBTc4 cells at OD, CD, and SD loci. Each row represents one individual locus. (B-C) Average RPKM-normalized BRG1 ChIP-seq signal 2 kb around OD, CD, and SD loci upon SOX2 depletion (B) and OCT4 depletion (C). (D-F) Frequency of overlap with motifs at OD, CD, and SD loci as well as in background regions (BG) for the canonical OCT4::SOX2 motif (D), the OCT motif (E), and the SOX motif (F). (G-I) Enrichment (-log(p)) values for the closest gene in the OD, CD, and SD groups in the gene ontology sets PluriNetWork (G), ESC Pluripotency Pathways (H), and the KEGG gene set “Signaling pathways regulating pluripotency” (I). (J-K) Average RPKM-normalized OCT4 (J) and SOX2 (K) ChIP-seq signal 2 kb around OD, CD, and SD loci upon SOX2 depletion (J) and OCT4 depletion (K)

To understand which features distinguish OD, SD, and CD loci, we performed motif analysis on the underlying sequences. While both OD and CD loci were strongly enriched for the OCT4::SOX2 canonical motif and the OCT motif, SD loci were more enriched for the SOX motif (Fig. 2D-F and Table 1). SD sites were also enriched for the AP-2 motif (Fig. S4B). TFAP2C, a member of the AP-2 family, is known to regulate differentiation into trophoblast stem (TS) cells together with SOX2 (Adachi et al., 2013). Interestingly, when reanalyzing data from TS cells (Adachi et al., 2013; Ishiuchi et al., 2019) we found SD sites to be highly SOX2-bound and accessible compared to OD and CD loci (Fig. S4C-D). Furthermore, SD loci were enriched near genes that increased in nascent mRNA expression upon loss of OCT4 (King and Klose, 2017) (Fig. S4E), which by itself leads to TS cell differentiation (Adachi et al., 2013). In contrast, OD and CD loci were enriched near genes that decreased in nascent mRNA expression upon OCT4 depletion (Fig. S4E). We next aimed to determine the fraction of pluripotency-associated enhancers falling in the OD, SD, and CD groups. To this end, we checked for enrichment of the nearest gene in three gene ontology (GO) sets associated specifically with pluripotency. We found that OD loci were most enriched in all three GO sets (Fig. 2G-I). We also analyzed the binding profiles of other pluripotency TFs (ESRRB, NANOG, KLF4, SALL4) (Aksoy et al., 2014; Chronis et al., 2017; Kim et al., 2018; Xiong et al., 2016) and found an enrichment in the CD group, although all these TFs bound to some extent to all groups of loci (Fig. S4F). Notably, all groups were also enriched for the “cell differentiation” GO term (Fig. S4G), in line with the role of OCT4 and SOX2 in ES cell differentiation. Since SOX2 was shown to require PARP1 to bind to a subset of genomic regions in ES cells (Liu and Kraus, 2017), we asked whether PARP1 dependence could explain the differential regulation of chromatin accessibility between these groups. We thus reanalyzed data from wt and PARP1 knockout (KO) ES cells (Gao et al., 2009; Yang et al., 2004), and found a reduction of SOX2 binding in PARP1 KO cells at OD, CD, and SD loci (Fig. S4H). Thus, PARP1 dependence cannot explain the differential regulation of chromatin accessibility between OD, CD, and SD loci. Overall, these results indicate that OCT4 and SOX2 regulate partially independent sets of pluripotency and differentiation enhancers, with OCT4 having the largest influence on chromatin accessibility of pluripotency-associated regulatory elements.

**Table 1.**
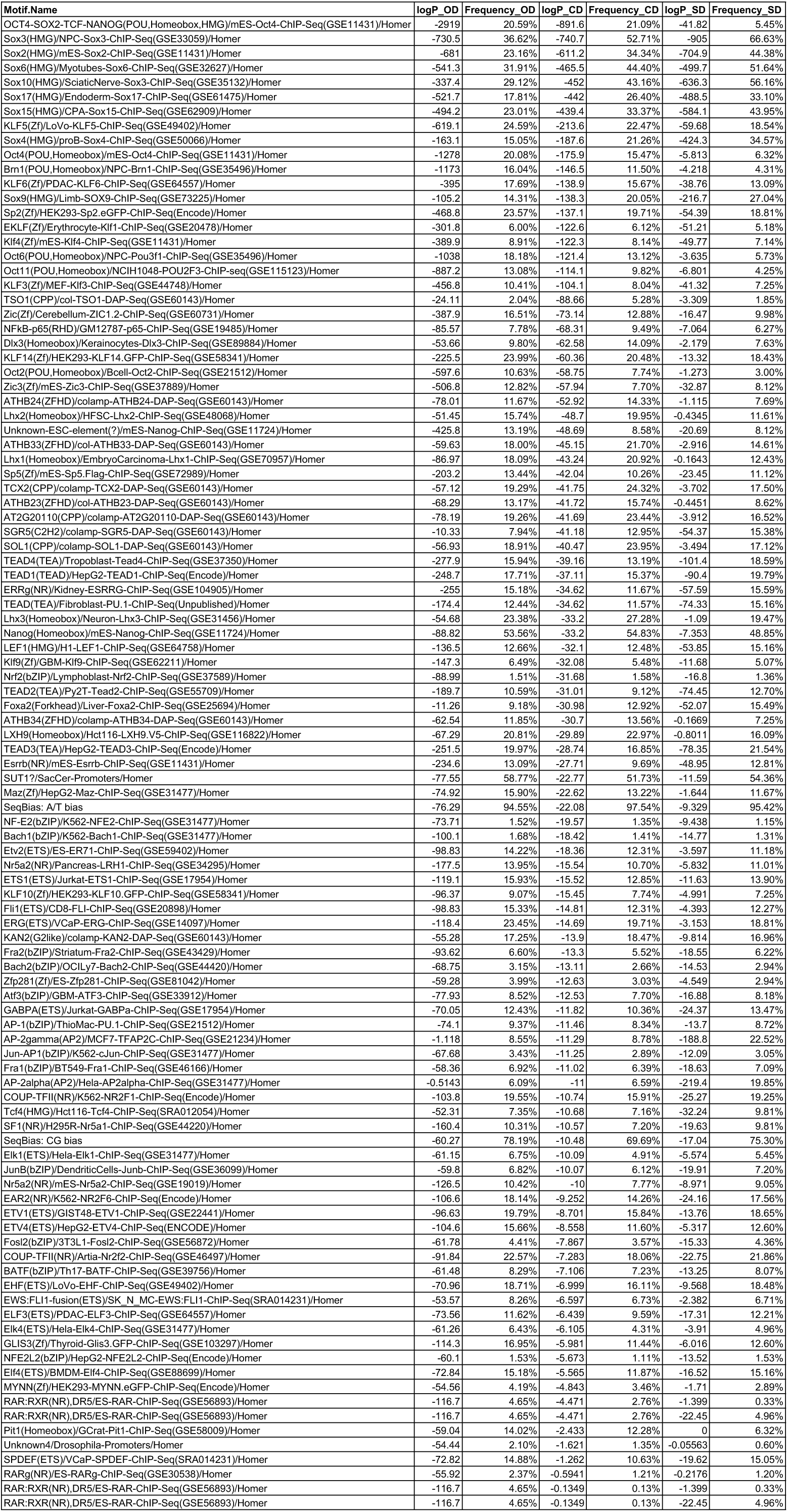
Enrichment (logP) and frequency of known motifs from HOMER in the sequences of OD, CD, and SD loci.

### Cooperative binding between OCT4 and SOX2 is mainly mediated indirectly through changes in chromatin accessibility

Several lines of evidence suggest that OCT4 and SOX2 exhibit cooperative DNA binding. In vitro electrophoretic mobility shift assays and fluorescence correlation spectroscopy experiments have shown that OCT4 and SOX2 display enhanced binding to the OCT4::SOX2 motif when binding together (Mistri et al., 2015, 2018). While in vitro experiments reported OCT4-assisted binding on a purified nucleosomal template (Li et al., 2019), single-molecule imaging in live ES cells (Chen et al., 2014) and ChIP-seq analysis of OCT4 in the presence or absence of SOX2 in fibroblasts (Raccaud et al., 2019) have provided evidence that SOX2 assists OCT4 binding in vivo. However, while these experiments suggest that OCT4 and SOX2 can display direct cooperativity, the role this mechanism plays in their colocalization in the complex in vivo chromatin and nuclear environment is unclear. We reasoned that the independent regulation of chromatin accessibility by OCT4 and SOX2 at a large number of loci could result in indirect cooperativity, i.e. each TF could assist the binding of the other through increasing chromatin accessibility. In line with this hypothesis, it was previously shown that upon loss of OCT4, SOX2 binding loss is correlated to the loss in chromatin accessibility (King and Klose, 2017). However, since most of the in vivo evidence points at a role for SOX2 in mediating cooperative OCT4 DNA-binding rather than vice versa (Chen et al., 2014; Raccaud et al., 2019), we interrogated genome-wide binding of OCT4 upon loss of SOX2 using ChIP-seq in 2TS22C cells treated with dox for 26 hours. We found that changes in OCT4 binding were also highly correlated to changes in chromatin accessibility upon SOX2 loss (Pearson’s R=0.77) (Fig. S5A). We next analyzed OCT4 and SOX2 binding in the presence or absence of SOX2 and OCT4, respectively, at OD, CD, and SD loci. We found that OCT4 binding was only slightly decreased at OD sites in the absence of SOX2, while SOX2 binding at SD sites was mildly increased in the absence of OCT4 (Fig. 2J-K). These findings were also consistent when narrowing down our analysis to sites containing a canonical OCT4::SOX2 motif (Fig. S5B-E). The slight loss of OCT4 binding at OD sites in the absence of changes in accessibility suggests that other mechanisms such as recruitment by SOX2 may play a role in the binding of OCT4, in line with SOX2 enhancing OCT4 binding.

Upon loss of its partner protein, OCT4 loses binding at 10’264 loci and gains binding at 1’153 loci, while SOX2 loses binding at 7’610 loci and gains binding at 4’423 loci. This indicates that the ability of OCT4 to occupy its specific binding sites is more impacted by the absence of SOX2 than vice-versa, and that SOX2 can get rerouted to new loci in the absence of OCT4. We further noticed that loci gaining accessibility upon loss of OCT4, which are highly enriched for differentiation terms (Fig. S4G), also gained binding by SOX2 (see Fig. S2A columns 6-7) and were enriched for the SOX and AP-2 motifs (Table 2). 3’484 loci displayed a significant increase in both accessibility and SOX2 binding. Interestingly, these loci decreased their accessibility upon SOX2 loss (Fig. S5F) and gained BRG1 occupancy concomitantly with OCT4 loss (Fig. S5G), in line with SOX2 recruiting the BAF complex to promote chromatin opening. This may suggest that OCT4 sequesters SOX2 to OCT4-SOX2 sites, and upon OCT4 loss SOX2 is free to bind and increase the accessibility of differentiation-associated regulatory elements. Overall, these results indicate that cooperative binding of OCT4 and SOX2 in ES cells is mainly mediated indirectly through changes in chromatin accessibility. However, while SOX2 enhances OCT4 binding in general, the presence of OCT4 reroutes SOX2 to pluripotency loci.

**Table 2.**
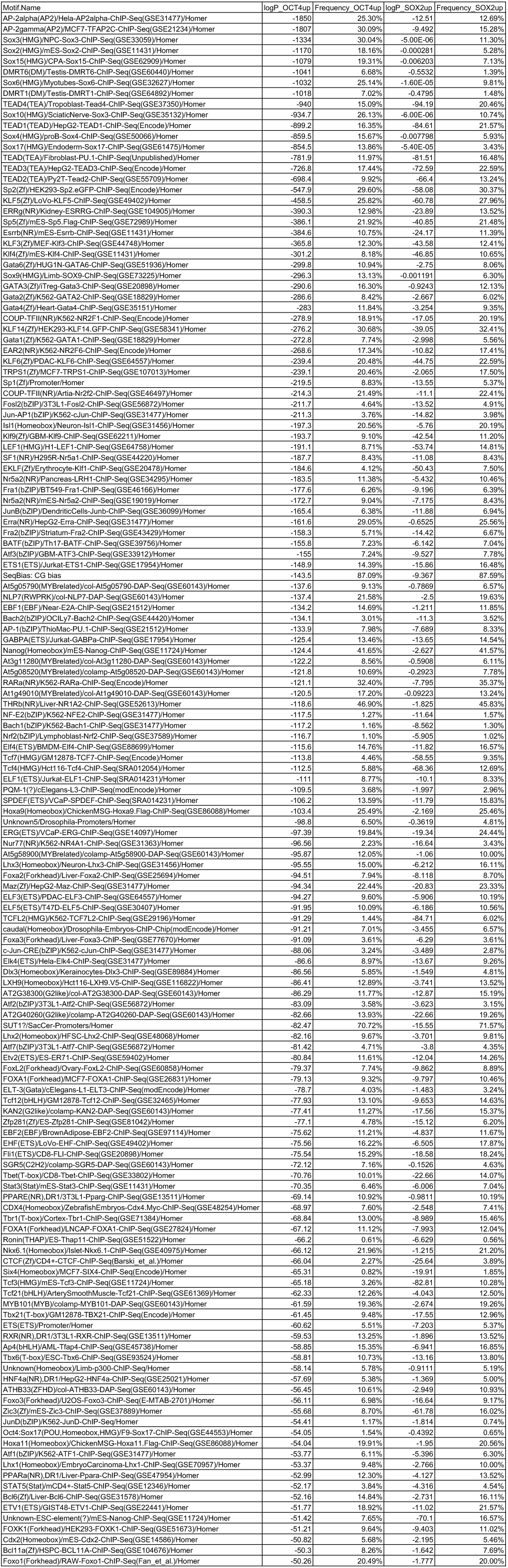
Enrichment (logP) and frequency of known motifs from HOMER in the sequences of loci gaining accessibility upon loss of OCT4 (OCT4up) and SOX2 (SOX2up).

### OCT4 is required at the M-G1 transition to re-establish enhancer accessibility

Transient depletion of OCT4 or SOX2 at the M-G1 transition has been shown to hinder pluripotency maintenance (Liu et al., 2017; Deluz et al., 2016), but the underlying mechanisms are not known. This time window coincides with enhancer reopening upon chromatin decompaction, but whether pioneer factors are involved in this process is not clear. As we found OCT4 to have the broadest influence on accessibility of pluripotency-associated loci, we focused on its role in regulating chromatin accessibility at the M-G1 transition. To allow near-complete loss of OCT4 at the M-G1 transition, we used ZHBTc4 cells constitutively expressing OCT4 fused to a SNAP-tag and a Cyclin B1 degron (mitotic degron; MD) or a mutated version thereof (MD*; Fig. 3A), which have been described previously (Kadauke et al., 2012). Importantly, lower than wildtype levels of OCT4 have been shown to sustain or even enhance pluripotency (Karwacki-Neisius et al., 2013; Radzisheuskaya et al., 2013). We thus reasoned that OCT4 levels need to decrease below a certain threshold to impact chromatin accessibility of pluripotency regulatory elements. Furthermore, the MD strategy strongly decreases but does not fully eliminate the target protein (Deluz et al., 2016; Liu et al., 2017). We therefore expressed MD-OCT4 and MD*-OCT4 at lower than wildtype levels from the constitutively active but relatively weak PGK promoter. After lentiviral transduction of the constructs, we stained cells with the SNAP-Cell 647-SiR dye (Lukinavičius et al., 2013) and sorted for a narrow window of SNAP expression to obtain the same average level of OCT4 tagged with MD and MD* across the cell cycle, as described previously (Deluz et al., 2016) (Fig. S6A). We also transduced cells to express YPet-MD in a constitutive manner, which allows for discrimination between cells in early G1 (YPet-negative) and late G1 phase (YPet-positive). In combination with Hoechst staining, this enables sorting cells in early G1 (EG1), late G1 (LG1), S, and late S/G2 (SG2) phase as described previously (Kadauke et al., 2012) (Fig. S6B). SNAP-MD-OCT4 levels were correlated with YPet-MD levels in flow cytometry, indicating that OCT4 levels are restored in LG1 in MD-OCT4 cells (Fig. S6C), as shown previously (Liu et al., 2017). In the absence of dox, these cell lines display no substantial difference in chromatin accessibility at OCT4-regulated loci (Fig. S6D). When grown in the presence of dox, MD*-OCT4 cells maintain a higher fraction of dome-shaped colonies, higher alkaline phosphatase activity, higher expression of pluripotency markers and lower expression of differentiation markers (Fig. S6E-G) than MD-OCT4 cells, as also shown previously (Liu et al., 2017).

To test whether depletion of OCT4 at the M-G1 transition affects chromatin accessibility, we treated cells with dox for 40 hours to ensure that all cells have gone through at least one cell division expressing only MD or MD*-tagged OCT4. Note that dox-treated cells had a longer G1 phase as compared to wt ES cells, as shown before to be a consequence of lower than wt OCT4 levels (Lee et al., 2010). However, there was no difference in the proportion of cells in the different cell cycle phases between MD-OCT4 and MD*-OCT4 (Fig. S6H). We sorted cells in EG1, LG1, S, and SG2 phases, performed ATAC-seq, and compared the accessibility between MD-OCT4 and MD*-OCT4 cells at each cell cycle phase. OCT4-regulated loci that increased or decreased in accessibility upon complete OCT4 depletion (see Fig. 1B) were also affected by transient M-G1 degradation (Fig. S6I-J, Fig. 3B-C). This shows that OCT4 is required at the M-G1 transition to restore chromatin accessibility and that loci gaining accessibility upon OCT4 loss are also dynamically regulated by OCT4 levels.

To characterize the different dynamic behaviors of chromatin accessibility changes across the cell cycle, we used k-means clustering on the change in accessibility between MD-OCT4 and MD*-OCT4 cells at all accessible loci displaying an OCT4 ChIP-seq peak (Fig. 3D). Two clusters displayed decreased accessibility in EG1 and recovered their accessibility incompletely (cluster 1) or completely (cluster 2) over the cell cycle. Notably, cluster 2 loci were less affected in EG1 than cluster 1 loci, which likely explains their complete recovery. Cluster 3 loci were characterized by a minor decrease in accessibility but that persisted over the cell cycle, and cluster 4 loci were unaffected by OCT4 loss. In contrast to clusters 1-3, cluster 4 was enriched near TSSs (Fig. S7A) and for the H3K4me3 promoter mark (Cao et al., 2018) (Fig. S7B), in line with OCT4 generally not affecting accessibility at promoters (Fig.1B and (King and Klose, 2017)). However, cluster 4 also contained many loci enriched for active enhancer marks (H3K4me1 and H3K27ac) (Kumar et al., 2016; Rickels et al., 2017), similar to clusters 1-3 (Fig. S7B). To test whether active histone marks also acutely change upon rapid loss of OCT4, we performed ChIP-seq for H3K27ac across the cell cycle in cells expressing MD-OCT4 or MD*-OCT4. The difference in H3K27ac across the cell cycle between the SNAP-MD-OCT4 and SNAP-MD*-OCT4 cell lines mimicked the corresponding changes in accessibility, although with smaller amplitude (Fig. S7C-D), suggesting that this modification is also highly dynamic and sensitive to OCT4 levels. We analyzed the fraction of regions in the different clusters overlapping previously annotated ES cell super-enhancers (SEs) and “typical” enhancers (TEs) (Sabari et al., 2018). We found these to be enriched in all clusters compared to non-OCT4 bound regions, with slightly more enrichment in clusters 1 and 3 for both SEs and TEs. This suggests that a large fraction of both SEs and TEs are permanently affected by the transient loss of OCT4 at the M-G1 transition (Fig. S7E).

As mentioned before, pluripotency was shown to be maintained at lower than wild type OCT4 expression levels. To ask whether chromatin accessibility of the observed clusters was OCT4 level-dependent within a higher OCT4 concentration range, we interrogated chromatin accessibility in the context of physiological variations of OCT4 levels. To do so, we took advantage of ATAC-seq data we previously acquired on cells differing in their OCT4 levels by a factor of ∼2, due to temporal fluctuations in their endogenous levels (Strebinger et al., 2019). We compared chromatin accessibility of the different clusters we identified for cells expressing high versus low endogenous levels of OCT4 and found virtually no differences in chromatin accessibility between these groups across all clusters (Fig.S7F), consistent with the ability of moderately low OCT4 levels to fully sustain pluripotency.

To understand the reason for the differential impact of transient OCT4 depletion on chromatin accessibility, we performed motif search analysis and compared OCT4 binding at the different clusters. We found a higher enrichment for the canonical OCT4::SOX2 motif (Fig.3E) and a higher OCT4 occupancy (Fig.3F) at cluster 1 loci. Consistently, cluster 1 contained mostly OD and CD loci identified above (Fig. S7G). We did not find strong differential enrichment for other motifs that could explain the differential regulation of these loci (Table 3). As high OCT4 binding was a signature of the loci most sensitive to transient OCT4 loss, we next aimed to determine the relationship between OCT4 binding and chromatin accessibility. We compared chromatin accessibility in ZHBTc4 cells in the presence or absence of OCT4 in conditions with matched OCT4 ChIP-seq and ATAC-seq data (King and Klose, 2017). The OCT4 ChIP-seq signal was correlated to loss of accessibility upon OCT4 depletion (Fig. S8A) as shown previously, but also to chromatin accessibility in untreated cells (Fig. S8B), indicating that strong OCT4 binding sites are both highly accessible and sensitive to OCT4 levels. Taken together, these results reveal different classes of OCT4-bound loci that show different cell cycle accessibility dynamics upon OCT4 loss at the M-G1 transition, and that highly bound sites are particularly accessible and sensitive to OCT4 loss for the maintenance of their accessibility and H3K27 acetylation.

**Table 3.**
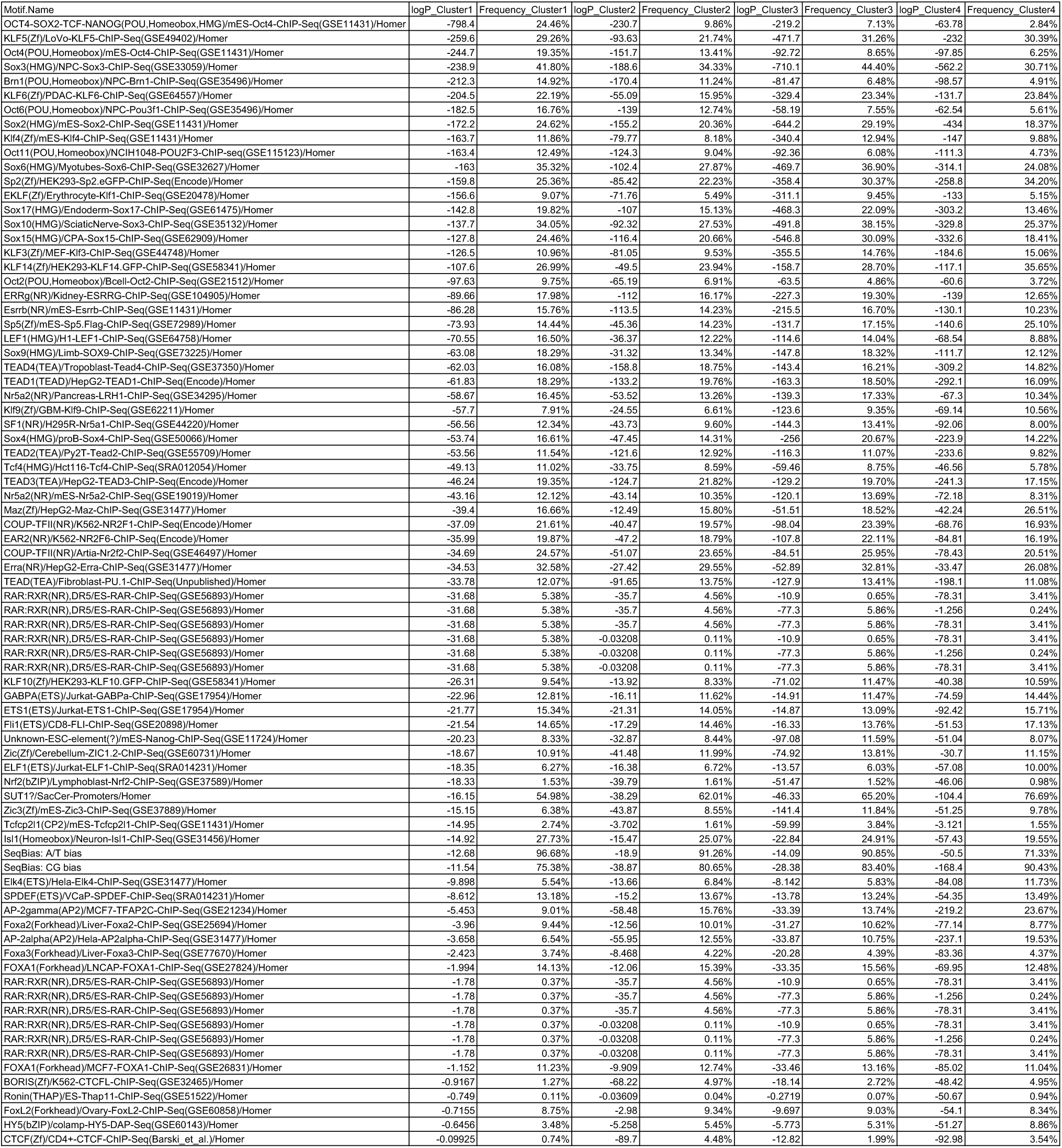
Enrichment (logP) and frequency of known motifs from HOMER in the sequences of the different clusters in Fig. 3D.

### OCT4 is required throughout the cell cycle to maintain enhancer accessibility

We next asked whether OCT4 also plays a role in maintaining enhancer accessibility in other cell cycle phases. To do so, we generated a cell line allowing drug-inducible degradation of OCT4. Briefly, we used lentiviral vectors to constitutively express the Tir1 ubiquitin ligase (allowing Auxin-inducible ubiquitination and degradation of target proteins (Dharmasiri et al., 2005; Kepinski and Leyser, 2005)) and OCT4 fused to mCherry and an Auxin-inducible degron tag (Morawska and Ulrich, 2013; Nishimura et al., 2009) (mCherry-OCT4-AID) in ZHBTc4 cells (Fig. 4A). To ensure near-complete OCT4 depletion, we expressed OCT4-AID at low levels using the PGK promoter. We further expressed YPet-MD in this cell line to allow for cell sorting in different cell cycle phases, as described above. Upon addition of indole-3-acetic acid (IAA, also known as Auxin), the AID-tagged OCT4 displayed an exponential degradation profile with a half-life of 0.32 h (Fig. 4B). After IAA washout, OCT4 recovered to approximately half of the concentration before IAA treatment within ∼4.5 hours (Fig.4C), in line with the OCT4 protein half-life of ∼4 hours (Alber et al., 2018).

**Figure 4.**
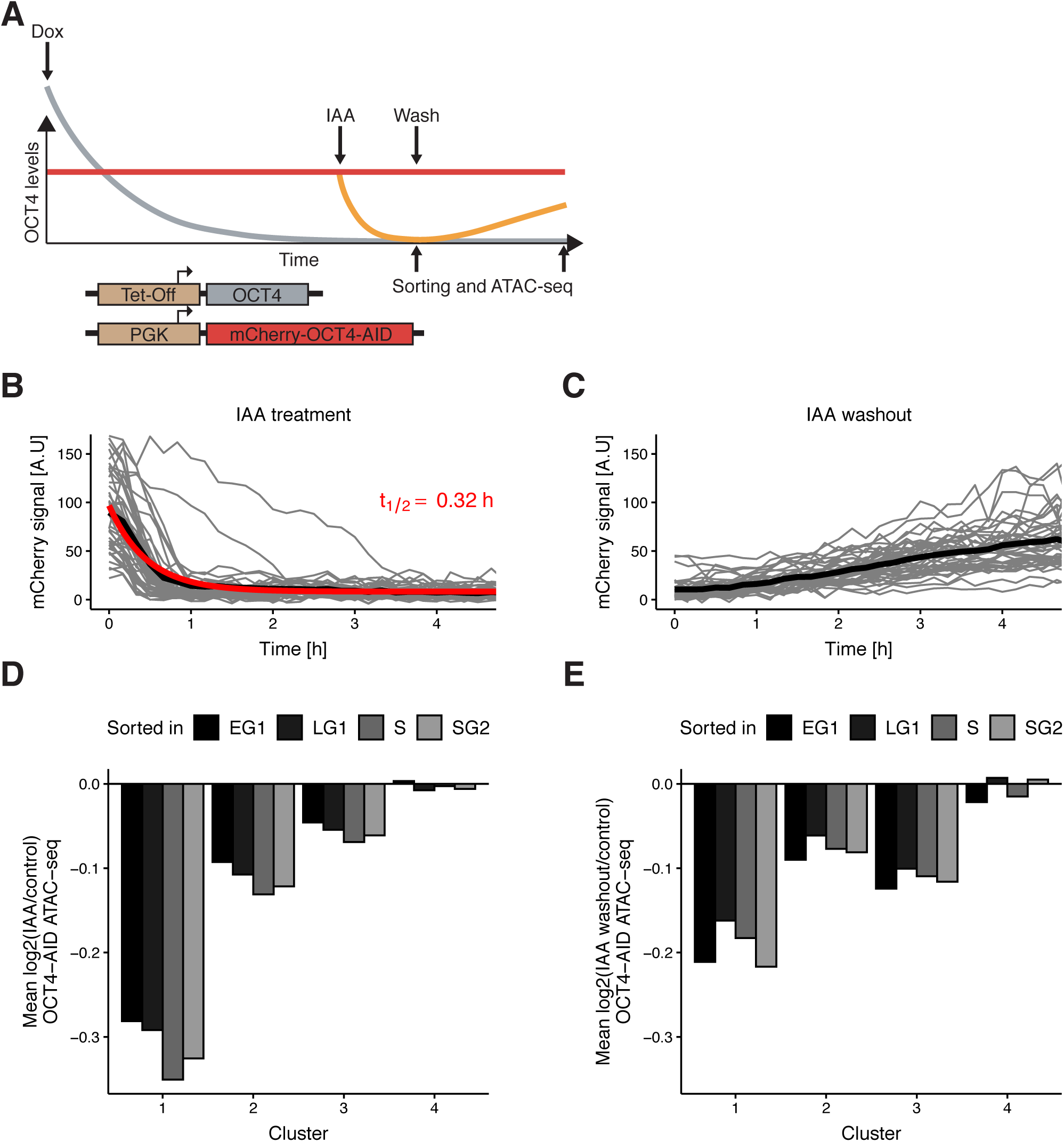
Auxin-inducible degradation reveals pioneer activity of OCT4 at different cell cycle phases. (A) Experimental strategy used to assess the impact of OCT4 depletion and recovery at different cell cycle phases. (B) Red fluorescence (mCherry) signal in mCherry-OCT4-AID cells treated with IAA at t=0 as measured by fluorescence microscopy. Gray lines: single cell traces; Black line: population average; Red line: exponential fit. Red text: half-life value derived from the exponential fit. (C) Red fluorescence (mCherry) signal in mCherry-OCT4-AID treated with IAA for 2.5h and then washed out at t=0 as measured by fluorescence microscopy. Gray lines: single cell traces; Black line: population average. (D) Average log2 fold-change values of accessibility between IAA-treated and untreated OCT4-AID cells in the four clusters from Fig. 3D at each cell cycle phase. (E) Average log2 fold-change values of accessibility between cells first treated with IAA and then washed out, compared to untreated OCT4-AID cells for the four clusters from Fig. 3D at each cell cycle phase.

To verify that OCT4 degradation kinetics are similar across the cell cycle, we applied IAA for 0.5 h (partial degradation) and 2 h (full degradation) before analyzing the mCherry signal by flow cytometry. At 2 hours of treatment, mCherry levels were similar to those of mCherry negative cells (Fig. S9A). We observed highly similar changes in the mCherry signal across all cell cycle phases (Fig. S9B-C), consistent with previous reports on the cell cycle-independence of Auxin-mediated protein degradation (Holland et al., 2012). OCT4 recovery after IAA washout was also very similar across the cell cycle (Fig. S9D). After addition of dox for 24 hours to remove untagged OCT4, we treated cells with IAA or not for 2 hours, sorted for different cell cycle phases, and performed ATAC-seq (Fig. 4A). The relative magnitude of change in accessibility in the different clusters was consistent with our mitotic degradation experiment (Fig. 4D). Remarkably, the average loss of accessibility was very similar at all cell cycle phases in clusters 1-3 (Fig. 4D, Fig. S9E).

Next, we quantified the recovery of chromatin accessibility across the cell cycle. We treated OCT4-AID cells with dox for 24 hours, then with IAA or not for 2.5h, washed out the drug and incubated cells for 4.5 hours, sorted cells in different cell cycle phases and performed ATAC-seq (Fig. 4A). While both cluster 1 (Fig. S9F) and 2 substantially recovered chromatin accessibility, cluster 3 loci did not (Fig.4E), in line with their permanent decrease of accessibility over the cell cycle upon OCT4 degradation at the M-G1 transition (see Fig. 3D). Overall, these data show that OCT4 is required across the cell cycle to maintain chromatin accessibility at enhancer elements.

### Dynamic relationship between OCT4 concentration and chromatin accessibility

We next aimed to quantify the dynamics of chromatin accessibility changes in response to OCT4 loss. Since residence times of OCT4 on specific DNA sites are in the second-range (Chen et al., 2014; Teves et al., 2016; Deluz et al., 2016), we reasoned that if continuous OCT4 re-binding is required to maintain chromatin accessibility, changes in chromatin accessibility and OCT4 concentration should occur in a quasi-synchronized manner. To test this hypothesis, we performed a time-course experiment by treating OCT4-AID cells with IAA for 0.5h, 1h, 2h, 3h, 4h, 6h, and 10h, and performed ATAC-seq at each time point. We took advantage of our different clusters, which showed differential response to OCT4 loss at the M-G1 transition (see Fig. 3D), and analyzed accessibility loss at these loci over time. At all OCT4-responsive clusters (1-3), accessibility loss was near-complete after 1 hour of IAA treatment (Fig. 5A-B), in line with accessibility being highly dynamic with OCT4 levels. At 6 and 10 hours of treatment, cluster 4 sites that were insensitive to OCT4 degradation at the M-G1 transition started to lose accessibility, suggesting a broader and potentially indirect impact of OCT4 loss on chromatin accessibility (Fig. 5A). We thus focused on the first 4 hours of OCT4 removal to estimate the kinetics of accessibility loss. We fitted a single-component exponential function including an offset to account for the residual ATAC-seq signal after OCT4 loss. At clusters 1-3, the half-life of accessibility loss was remarkably close to the half-life of OCT4-AID upon IAA treatment, i.e. around 0.5 hours (Fig. 5C-E). We were unable to fit an exponential decay to cluster 4, as expected from its OCT4-independent chromatin accessibility regulation (Fig. 5F). In summary, these data suggest that regulation of enhancer accessibility is extremely dynamic and requires the constant presence of OCT4.

**Figure 5.**
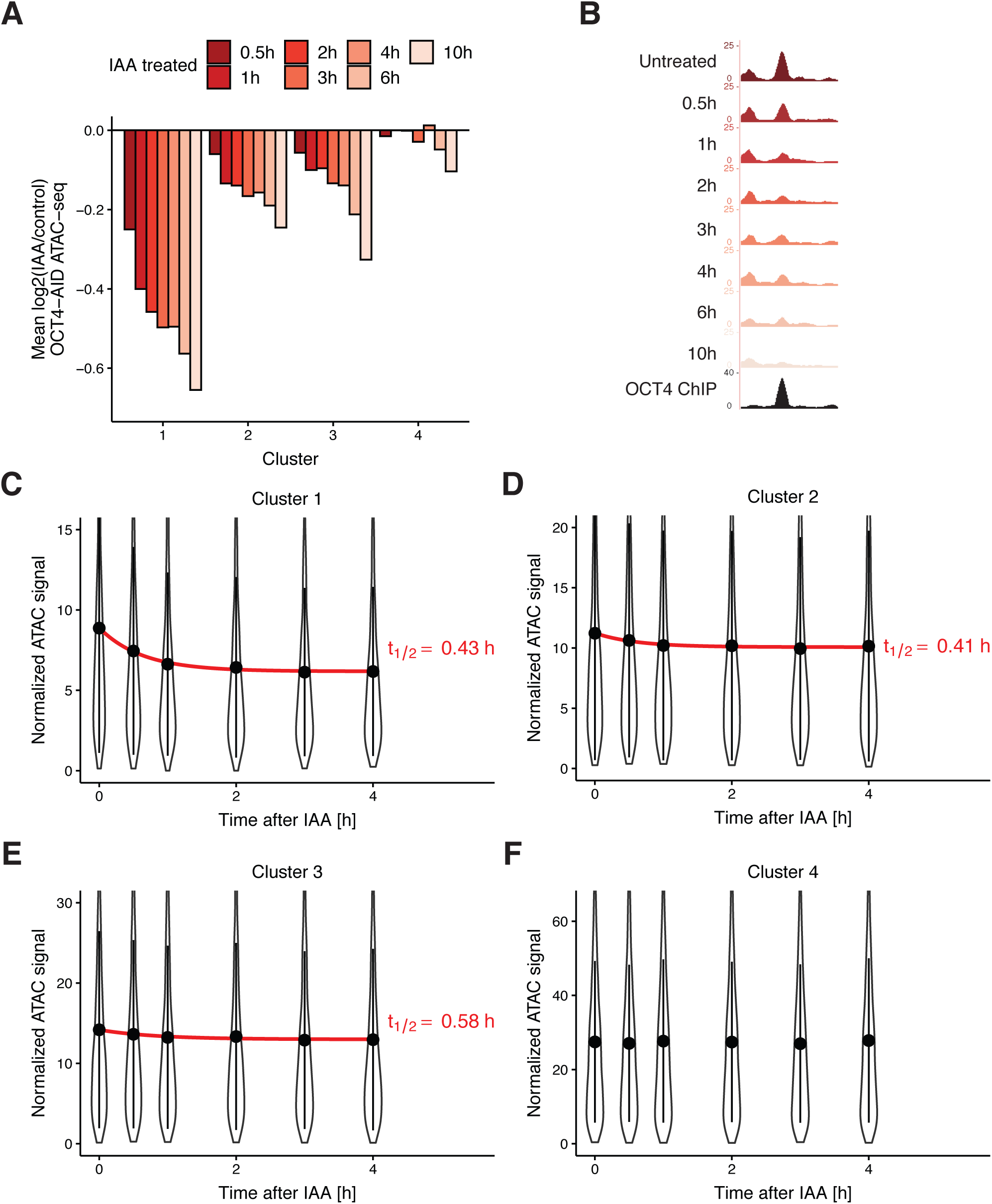
Time course analysis of chromatin accessibility changes during OCT4 degradation reveals its highly dynamic pioneer activity. (A) log2 fold-change values of accessibility compared to untreated cells in the four clusters from Fig. 3D at different time points of IAA treatment. (B) Genome browser track of accessibility profiles upon treatment with IAA for different durations at a cluster 1 locus at chr3:137779908-137780687. (C-F) Violin plot of normalized ATAC-seq signal across different time points in cluster 1 (C), cluster 2 (D), cluster 3 (E), and cluster 4 (F). Dots: mean; Vertical lines: standard deviation; Red lines in C-E: exponential fit; Red text in C-E: half-life value derived from the exponential fit.

## Discussion

In this study we dissected the roles and interplay of OCT4 and SOX2 in regulating chromatin accessibility in ES cells. To our surprise, we found a large number of enhancers that were bound by both transcription factors but for which chromatin accessibility was regulated by only one of them. In the future it will be interesting to explore whether differences in the topology of OCT4 and SOX2 binding sites on the nucleosome surface or genomic location-dependent DNA residence times could explain these findings. Our results also show that both OCT4 and SOX2 regulate the genomic occupancy of each other mainly via regulation of chromatin accessibility. This is reminiscent of dynamic assisted loading, in which two TFs assist the loading of each other to either the same or nearby DNA binding sites (Swinstead et al., 2016; Goldstein et al., 2017).

Surprisingly, upon OCT4 loss chromatin accessibility increased at a large number of genomic sites enriched for proximity to differentiation genes, even when OCT4 was degraded for a brief period of time at the M-G1 transition. The fact that SOX2 occupies these sites and is required to maintain their accessibility suggests that in the absence of OCT4, SOX2 is rerouted to these loci and promotes differentiation together with other partners such as TFAP2C. Therefore, the rapid action of OCT4 in early G1 phase might be required to ensure both the maintenance of chromatin accessibility at pluripotency enhancers and to silence differentiation enhancers. Whether the previously shown association of OCT4 to mitotic chromosomes (Deluz et al., 2016; Liu et al., 2017; Teves et al., 2016) facilitates its action in early G1 will require further investigation.

We found that OCT4 degradation leads to a rapid decrease in chromatin accessibility at all clusters of OCT4-regulated enhancers across the cell cycle with very similar kinetics, which tightly mirrored changes in OCT4 concentration and thus suggests highly dynamic regulation of chromatin accessibility by OCT4. However, the recovery of chromatin accessibility upon increase of OCT4 concentration displayed locus-dependent behavior. In contrast to clusters 1 and 2, cluster 3 loci did not recover over the time course of several hours either after M-G1 or auxin-induced degradation. While the mechanisms underlying these findings are unclear, permanent loss (cluster 3) or incomplete recovery (cluster 1) of chromatin accessibility may explain why OCT4 loss at the M-G1 transition results in impaired pluripotency maintenance.

Protein depletion by degron systems works by increasing protein degradation rates without affecting their synthesis rate. Therefore, they suffer from an inherent tradeoff in maximizing expression levels when the degron is inactive while minimizing residual expression level when the degron is active. Here we expressed OCT4 at relatively low levels to ensure sufficient depletion, allowing us to show that the pioneering function of OCT4 is required constantly and throughout the cell cycle to maintain enhancer accessibility. However, the relatively low dynamic range of accessibility changes prohibits sensitive detection of specific loci that are quantitatively more or less sensitive to OCT4 loss at different cell cycle phases. Furthermore, whether recurrent, transient loss of OCT4 outside of the M-G1 transition would also lead to pluripotency loss would have to be addressed in future studies. Here we found that OCT4 is constantly required to maintain chromatin accessibility in self-renewing ES cells. This is reminiscent of a recent study showing that the pioneer factor Zelda is required throughout zygotic genome activation in Drosophila for proper gene expression (McDaniel et al., 2019). In contrast, in the context of pituitary lineage specification PAX7 requires 72 hours to fully open melanotrope-specific enhancers but is subsequently not required to maintain these (Mayran et al., 2018). It is possible that PAX7 hands over the role of maintaining accessibility to other factors, such as TPIT (Mayran et al., 2019), and is only required at the transition between cell fates. This indicates that pioneering activity can have different manifestations that depend heavily on other regulatory factors and chromatin features. Pluripotent stem cells may be particularly dynamic in this regard, as they need to be able to quickly rewire their gene expression programs upon receiving differentiation signals. In contrast, more differentiated cell types may have mechanisms to avoid precocious changes in gene expression upon subtle alterations in the concentration of TFs. Therefore, the high sensitivity of enhancers to the concentration or activity of pioneer TFs in ES cells could serve as a mechanism to regulate cell fate with precise temporal control.

## Acknowledgments

This work was supported by the Swiss National Science Foundation (grants #PP00P3_179068 and PP00P3_17205 to D.M.S.). A.C.A.M.F. was supported by a Marie Curie Intra European Fellowship within the 7th European Community Framework Programme. This work was further supported by AgingX (SystemsX.ch) and SNF (310030_182655). We thank Bastien Mangeat, Elisa Cora, Paolo Ferrari, and Lionel Ponsonnet from the Gene Expression Core Facility for high-throughput sequencing, Miguel Garcia, Loïc Tauzin, Valérie Glutz, and André Mozes from the Flow Cytometry Core Facility for cell sorting, Olivier Burri and Romain Guiet from the Bioimaging and Optics Platform for assistance with cell tracking, the staff at Vital-IT and SCITAS for cluster computing, and Armelle Tollenaere for critical reading of the manuscript.

## Author contributions

D.M.S. and E.T.F. conceptualized the study and wrote the manuscript. E.T.F. performed the computational analysis. E.T.F., A.C.A.M.F., and C.D. performed ATAC-seq experiments. C.D. and E.T.F. performed ChIP-seq experiments. C.D. performed pluripotency assays, qRT-PCR, and immunofluorescence experiments. C.D. and S.G. generated cell lines. V.G. consulted on the bioinformatic analysis. D.M.S. and B.D. provided funding and resources.

## Data availability

All data are available upon demand. Sequencing data will be uploaded publicly upon publication.

## Methods

### Cell culture

Mouse ES cells were routinely cultured on cell culture-treated dishes coated with 0.1% gelatin (Sigma #G9391-100G) using the following culture medium: GMEM (Sigma #G5154-500ML) containing 10% ES-cell qualified fetal bovine serum (Gibco #16141-079), nonessential amino acids (Gibco #11140-050), 2mM L-glutamine (Gibco #25030-024), sodium pyruvate (Sigma #S8636-100ML), 100μM 2-mercaptoethanol (Sigma #63689-25ML-F), penicillin and streptomycin (BioConcept #4-01F00-H), in-house produced leukemia inhibitory factor (LIF), CHIR99021 (Merck #361559-5MG) at 3μM and PD184352 (Sigma #PZ0181-25MG) at 0.8μM. Cells were passaged by trypsinization (Sigma #T4049-100ML) every two to three days.

### Lentiviral vector production

Lentiviral vectors were produced by transfection of HEK 293T cells with the envelope (psPAX2, Addgene #12260), packaging (pMD2.G, Addgene #12259) (Dull et al., 1998), and lentiviral construct of interest using Calcium Phosphate transfection, as described previously (Suter et al., 2006). Viral vectors were concentrated 120-fold by ultracentrifugation at 20’000 rpm for 90 minutes at 4°C. 50’000 cells in 1 ml of medium in a 24-well plate were transduced with 50 μl of concentrated lentiviral vector particles to generate stable cell lines.

### Cloning of overexpression constructs

pLV-PGK-YPet-MD was derived from pLVTRE3G-Sox2-YPet-MD (Deluz et al., 2016) by amplification of YPet-MD and restriction cloning into pLV-rtTA3G-IRESBsd using AgeI and SalI. pLV-PGK-SNAP-MD-OCT4 and pLV-PGK-SNAP-MD*-OCT4 were derived by amplification of MD or MD* from SNAP-MD-SOX2 (Addgene #115687) and SNAP-MD*-SOX2 (Addgene #115688) (Deluz et al., 2016) and restriction cloning into pLVTRE3G-Oct4-YPet (Deluz et al., 2016) using SalI and XbaI. SNAP-MD-OCT4 and SNAP-MD*-OCT4 were further amplified and cloned by restriction cloning into pLV-rtTA3G-IRESBsd (Deluz et al., 2016) using AgeI and SalI. pLEX-mCherry-OCT4-AID was derived by amplification of OCT4 from pLV-PGK-SNAP-MD-OCT4, AID 71-114 from pEN244 (Addgene #92140) (Nora 2017), and mCherry from pLV-TRE3G-mCherry-PGK-Puro (Suter lab). mCherry and OCT4 were ligated using an XmaI restriction site and mCherry-OCT4 was ligated to AID using a KpnI restriction site. The mCherry-OCT4-AID fragment was cloned into the pLEX_305-C-dTAG backbone (Addgene #91798) (Nabet et al., 2018) using EcoRV and MluI restriction sites. pLV-pCAGGS-Tir1-V5 was derived by amplification of pCAGGS-Tir1-V5 from pEN395 (Addgene #92141) (Nora et al., 2017) and In-fusion cloning into pLV-PGK-SOX2-SNAP-IRES-Hygro (Strebinger et al., 2019) digested using XhoI and XbaI restriction enzymes.

### Generation of stable cell lines

To generate MD-OCT4 cell lines, ZHBTc4 cells were transduced with SNAP-MD-OCT4 and SNAP-MD*-OCT4 lentiviral vector particles and sorted to display near-identical average SNAP levels (Fig. S6A), subsequently transduced with PGK-YPet-MD lentiviral vector particles, and sorted to display near-identical average YPet-MD levels. To generate the OCT4-AID cell line, ZHBTc4 cells were transduced with pLV-pCAGGS-Tir1-V5 and PLEX-mCherry-OCT4-AID packaged lentiviral vectors (i.e Tir1-V5 and mCherry-Oct4-AID virus, respectively) and were selected with 2 μg/ml Puromycin (Thermo Fisher Scientific #A1113803) for 10 days. Subsequently, mCherry positive cells were sorted and transduced with PGK-YPet-MD lentiviral particles and sorted for YPet positive cells. Cells that displayed IAA-dependent degradation were selected by sorting a narrow window of mCherry-positive cells, followed by treatment with 500 nM IAA (Sigma #I5148-2G) for 1 hour, and sorting for mCherry-negative cells. All cell lines were maintained in medium without dox (Sigma #D3447-500MG) or IAA prior to experiments.

### Treatment conditions

ZHBTc4 YPet-MD SNAP-MD-OCT4 and SNAP-MD*-OCT4 were treated with 1 μg/ml dox for 40 hours prior to cell sorting. ZHBTc4 YPet-MD TIR1-V5 mCherry-OCT4-AID cells were treated with 1 μg/ml dox for 24 hours before adding IAA. Dox was maintained throughout the experiment. Cells were treated with 500 nM IAA (or not for control) for 2 hours or treated with 500 nM IAA (or not for control) for 2.5 hours, washed 5 times with PBS with 2 minutes incubation, and placed back in medium containing 1 μg/ml dox for 4.5 hours. For the time course experiment, OCT4-AID cells were seeded in different wells of a 24-well plate and treated with dox for 24 hours before adding IAA. Dox was maintained throughout the experiment. IAA was added at different time points (with one well left untreated) prior to cell collection. All wells were collected at the same time and subjected to ATAC-seq as described below.

### Cell cycle phase sorting

Cells were trypsinized, resuspended in culture medium with 50 μM Hoechst33342 (Thermo Fisher Scientific #H3570), and incubated for 15 minutes at 37°C. Cells were then spun down, resuspended in cold PBS with 1% FBS, and sorted according to their YPet-MD and Hoechst profile (Fig. S6B). Cells were sorted at 4°C into 1.5 ml Eppendorf tubes or 15 ml Falcon tubes containing a small amount of PBS with 1% FBS. Sorting for SNAP levels was done on a MoFlo Astrios (Beckman Coultier). All other sorting was done on a FACSAria II or a FACSAria Fusion (BD Biosciences).

### ATAC-seq

All ATAC-seq experiments were performed in biological duplicates. 50’000 cells were collected either directly after trypsinization or after sorting as described above and subjected to ATAC-seq as described previously (Buenrostro et al., 2013). All centrifugation steps were done at 800g at 4°C. Briefly, cells were centrifuged for 5 minutes and washed with cold PBS, then centrifuged for 5 minutes and resuspended in cold lysis buffer (10 mM Tris-HCl pH 7.4, 10 mM NaCl, 3 mM MgCl2, 0.1% NP-40), and centrifuged for 10 minutes. Subsequently, nuclei were resuspended in a solution of 0.5 μM Tn5 (in-house preparation according to (Chen et al., 2017)) in TAPS-DMF buffer (10 mM TAPS-NaOH, 5 mM Mgcl2, 10% DMF) and incubated for 30 minutes at 37°C. DNA was immediately purified using column purification (Zymo #D4004) and eluted in 10 μl nuclease-free H2O. Transposed DNA was amplified in a solution containing 1X NEBNextHigh Fidelity PCR Master Mix (NEB #M0541L), 0.5 μM of Ad1_noMX universal primer, 0.5 μM of Ad2.x indexing primer and 0.6x SYBR Green I (Thermo Fisher Scientific #S7585) using 72°C for 5 minutes, 98°C for 30 s, and 5 cycles of 98°C for 1s, 63°C for 30s, and 72°C for 60s. 10 μl of amplified DNA was analyzed by qPCR to determine the total number of cycles to avoid amplification saturation and accordingly amplified with additional 3-7 cycles of 98°C for 10s, 63°C for 30s, and 72°C for 60s. DNA was purified using column purification (Zymo #D4004) and size-selected by taking the unbound fraction of 0.55X AMPure XP beads (Beckman Coultier #A63880) followed by the bound fraction of 1.2X AMPure XP beads. Libraries were sequenced on an Illumina NextSeq 500 using 75 nucleotide read-length paired-end sequencing.

### ChIP-seq

All ChIP-seq experiments were performed in biological duplicates. Roughly 10 million cells per sample were collected after trypsinization and fixed with 2 mM disuccinimidyl glutarate (Thermo Fisher Scientific #20593) in PBS for 50 minutes at room temperature, spun down at 600g for 5 minutes and washed once with PBS. Cells were then treated with 1% formaldehyde (Axon Lab #A0877,0500) for 10 minutes at room temperature and quenched with 200mM Tris-HCl pH 8.0 for 10 minutes, washed with PBS and spun down. For ZHBTc4 YPet-MD SNAP-MD-OCT4 and SNAP-MD*-OCT4 cells, cells were subsequently resuspended in cold PBS with 1% FBS and at least 500’000 cells per cell cycle phase were sorted. Fixed cell pellets were kept on ice and resuspended in LB1 (50mM HEPES-KOH pH 7.4, 140mM NaCl, 1 mM EDTA, 0.5mM EGTA, 10% Glycerol, 0.5% NP-40, 0.25% Triton X-100), incubated 10 min at 4°C, spun down at 1700g, and resuspended in LB1 a second time, spun down and resuspended in LB2 (10mM Tris-HCl pH 8.0, 200mM NaCl, 1 mM EDTA, 0.5mM EGTA), incubated for 10 min at 4°C, spun down and washed without disturbing the pellet three times with SDS shearing buffer (10mM Tris-HCl pH 8.0, 1mM EDTA, 0.15% SDS) and finally resuspended in SDS shearing buffer. All buffers contained Protease Inhibtor Cocktail (Sigma #P8340-1ML) at 1:100 dilution. Chromatin was sonicated for 20 min at 5% duty cycle, 140 W, 200 cycles on a Covaris E220 focused ultrasonicator. Sonicated chromatin was equilibrated to 1% Triton X-100 and 150 mM NaCl and incubated with each antibody overnight at 4°C. Antibodies used were anti-BRG1 (Abcam #ab110641) at 5 μg per 10 million cells, anti-OCT4 (Cell Signaling Technology #5677S) at 20 μl per 10 million cells, and anti-H3K27ac (abcam #ab4729) at 2 μg/25 μg chromatin. Protein G Dynabeads (Thermo Fisher Scientific #10003D) were blocked with 5 mg/ml BSA in PBS, added to chromatin, and incubated at 4°C for 3 hours. Beads were washed twice with Low Salt wash buffer (10mM Tris-HCl pH 8.0, 150mM NaCl, 1 mM EDTA, 1% Triton X-100, 0.15% SDS, 1 mM PMSF), once with High Salt wash buffer (10mM Tris-HCl pH 8.0, 500mM NaCl, 1 mM EDTA, 1% Triton X-100, 0.15% SDS, 1 mM PMSF), once with LiCl wash buffer (10mM Tris-HCl pH 8.0, 1mM EDTA, 0.5mM EGTA, 250mM LiCl, 1% NP40, 1% sodium deoxycholate, 1mM PMSF), and finally with 1X TE (10 mM Tris pH 8.0, 1 mM EDTA) before being resuspended in ChIP Elution buffer (10 mM Tris pH 8.0, 1 mM EDTA, 1% SDS, 150 mM NaCl) with 400 ng/μl Proteinase K (Qiagen #19131) and reverse-crosslinked overnight at 65°C. DNA was purified using a MinElute PCR purification kit (Qiagen #28004) and libraries were prepared with the NEBNext Ultra II DNA Library Prep Kit (NEB #E7645S). Libraries were sequenced on an Illumina NextSeq 500 using 75-nucleotide read length paired-end sequencing.

### Pluripotency assays and qRT-PCR

ZHBTc4 YPET-MD cells expressing SNAP-MD-OCT4 or SNAP-MD*-OCT4 were plated at 400 cells per well in a 6-well plate with ES cell medium (see above) with 0 or 1 μg/ml dox and medium was refreshed every other day. At day 8, flat and dome-shaped colonies were scored according to morphology followed by alkaline phosphatase staining (Sigma #86R-1KT). For qRT-PCR experiments, cells were collected at day 8 and RNA was extracted using GenElute Mammalian Total RNA MiniPrep Kit (Sigma #RTN350). cDNA was synthesized using oligodT primers using SuperScript II Reverse Transcriptase (Thermo Fisher Scientific #18064014). qPCR was performed on a 7900HT (Applied Biosystems).

### Immunofluorescence microscopy

2TS22C and ZHBTc4 cells were plated in a 96-well plate coated for 1 hour at 37°C with 1:25 diluted StemAdhere (Primorigen Biosciences #S2071-500UG), treated with 1 μg/ml dox for different durations and fixed with 2% formaldehyde for 30 minutes at room temperature, washed with PBS, permeabilized with PBS with 5% FBS and 0.5% Triton X-100 for 30 minutes at room temperature, and incubated with the primary antibody, anti-OCT4 C-10 (Santa Cruz #sc-5279) at 1:500 dilution and anti-SOX2 (ThermoFisher #48-1400) at 1:200 dilution, in PBS with 5% FBS and 0.1% Triton X-100 at 4°C overnight. After washing with PBS, cells were incubated with the secondary antibody, anti-Mouse IgG AF488 (Thermo Fisher Scientific #A28175) and anti-Rabbit IgG AF647 (Thermo Fisher Scientific #A27040) at 1:1000, in PBS with 5% FBS and 0.1% Triton X-100 for 60 minutes at room temperature, with 2 ng/ml DAPI (Thermo Fisher Scientific #62248) for 10 minutes at room temperature and subsequently washed and imaged on an IN Cell Analyzer 2200 (GE Healthcare). Images were background-subtracted using FiJi (Schindelin et al., 2012) with a rolling ball radius of 50 pixels and analyzed using CellProfiler (Carpenter et al., 2006). Nuclei were identified using the Watershed module on the DAPI channel, objects that were too large or too small were discarded, and the integrated intensity in the OCT4 and SOX2 channels was measured within the identified nuclei.

### Time-lapse microscopy

ZHBTc4 YPet-MD TIR1-V5 mCherry-OCT4-AID cells were plated in a 96-well plate coated for 1 hour at 37°C with 1:25 diluted StemAdhere (Primorigen Biosciences #S2071-500UG) and imaged on an IN Cell Analyzer 2200 (GE Healthcare) using the TexasRed and Brightfield channels. Cells were treated with IAA or washed just prior to imaging. Images were background-subtracted using FiJi (Schindelin et al., 2012) with a rolling ball radius of 50 pixels and nuclei were tracked manually over time using the Manual Tracking plugin in FiJi in the Brightfield channel. The mean signal in 10 pixels around the tracked spot were measured in the TexasRed channel and the mean background signal at an equivalent sized spot free from cells (background) was subtracted at each time point.

### Data analysis for ATAC-seq and ChIP-seq

All sequencing libraries were aligned to the mm10 Mus musculus genome (GRCm38) with STAR 2.6.1c (Dobin et al., 2013) and duplicate reads were removed using Picard (Broad Institute). Reads not mapping to chromosomes 1-19, X, or Y were removed. Peaks were called with MACS 2.1.1.20160309 (Zhang et al., 2008) with settings ‘-f BAMPE -g mm’. For comparative analysis of ZHBTc4 and 2TS22C cells, all peaks from ZHBTc4 and 2TS22C ATAC-seq experiments were merged with BEDTools (Quinlan and Hall, 2010). For MD-OCT4 and OCT4-AID analyses, all peaks from ATAC-seq experiments in the corresponding cell lines were merged. For MD-OCT4 H3K27ac analysis, peak coordinates were expanded by 500 bp on both sides to account for the enrichment profile of H3K27ac. All peaks larger than 5 kb, overlapping peaks called in Input (no immunoprecipitation) samples from ES cells in S2iL (GSE89599) or SL (GSE87822), or overlapping blacklisted peaks (ENCODE Project Consortium, 2012) were removed. The HOMER2 (Heinz et al., 2010) functions makeTagDirectory and annotatePeaks.pl with settings ‘-noadj -len 0 -size given’ were used for read counting and count tables were loaded into RStudio. TMM Normalization was done with edgeR (Robinson et al., 2010) and analysis of differentially accessible regions was done with limma (Ritchie et al., 2015). Contrasts were designed as ∼0+Condition+Replicate, where Condition specifies the cell line and treatment and Replicate the date of the experiment, to take into account the paired nature of the experiments. For comparing unpaired experiments, i.e. untreated ZHBTc4 vs 2TS22C cell lines or untreated ZHBTc4 in SL versus S2iL, ∼0+Condition was used. For Fig. S8A-B, the mean of the TMM-normalized reads in the ChIP-seq and ATAC-seq replicates was divided by the nucleotide length of each region. For Figures 5C-F, the mean of the TMM-normalized reads in the replicates was used. The HOMER2 function findMotifsGenome.pl was used with the setting ‘-size given’ for motif searching. The most frequent known motif in target regions of a given class of known motifs (i.e. different versions of SOX and OCT motifs) was used. Background was calculated as the mean of HOMER-estimated background frequency in all groups/clusters. Only motifs with logP < -50 are shown in Tables 1-3. For GO analysis, the closest Entrez gene entry TSS to each region in groups was used and enrichment was calculated using the HOMER2 function findGO.pl with setting ‘mouse’. Gene names were converted between assemblies using biomaRt (Durinck et al., 2005). Replicate bam files were merged using SAMTools (Li et al., 2009) and converted to bigWig files using the deepTools 3.1.3 (Ramírez et al., 2016) function bamCoverage with settings ‘--normalizeUsingRPKM’. Average lineplots were generated using deepTools computeMatrix (with setting ‘reference-point’) and custom R code. Heatmaps were generated using the deepTools function plotHeatmap. Genome tracks were made in the UCSC genome browser (Kent et al., 2002). Plots were generated using ggplot2 (Wickham, 2009). Overlap between genomic regions was determined using GenomicRanges (Lawrence et al., 2013). The heatmap in Fig. 1C was generated using ComplexHeatmap (Gu et al., 2016). Color schemes were taken from colorbrewer2.org and https://rpubs.com/Koundy/71792.

### Published datasets

Published data in Table 4 were aligned and processed as described above. Processed bigWig files in Table 5 were downloaded from GEO (Edgar et al., 2002) or cistromeDB (Mei et al., 2017). When necessary, peak files were converted to mm9 using liftOver (Hinrichs et al., 2006). OCT4 and SOX2 ChIP-seq peaks were derived from newly generated (2TS22C OCT4) and published (ZHBTc4 OCT4 and SOX2) (King and Klose, 2017) datasets as described above as well as from processed SOX2 ChIP-seq peaks from asynchronous E14 cells (GSE89599) (Deluz et al., 2016) and merged with BEDTools (Quinlan and Hall, 2010). Super-enhancer and typical enhancers were taken from (Sabari et al., 2018) and converted to mm10 using liftOver (Hinrichs et al., 2006). ChromHMM tracks from mouse ES cells were downloaded from https://github.com/guifengwei/ChromHMM_mESC_mm10 (Pintacuda et al., 2017). ATAC-seq data from OCT4 high and OCT4 low sorted cells were taken from a previous study (Strebinger et al., 2019) and processed as described above, merging SHOH and SLOH samples into OCT4 high and SLOL and SHOL samples into OCT4 low.

**Table 4.** Description of publicly available data used in this study that was aligned and processed according to the Methods section.

| Experiment | Cell line | Expression construct | Treatment | Cell cycle phase | Antibody | Replicate | Accession | SRA | SRA | SRA | SRA |
| --- | --- | --- | --- | --- | --- | --- | --- | --- | --- | --- | --- |
| ATAC | ZHBTc4 | - | UNT | - | - | 1 | GSE87822 | SRR44137754 | SRR4413775 | SRR4413776 | SRR4413777 |
| ATAC | ZHBTc4 | - | DOX | - | - | 1 | GSE87822 | SRR4413786 | SRR4413787 | SRR4413788 | SRR4413789 |
| ATAC | ZHBTc4 | - | UNT | - | - | 2 | GSE87822 | SRR4413778 | SRR4413779 | SRR4413780 | SRR4413781 |
| ATAC | ZHBTc4 | - | DOX | - | - | 2 | GSE87822 | SRR4413790 | SRR4413791 | SRR4413792 | SRR4413793 |
| ATAC | ZHBTc4 | - | UNT | - | - | 3 | GSE87822 | SRR4413782 | SRR4413783 | SRR4413784 | SRR4413785 |
| ATAC | ZHBTc4 | - | DOX | - | - | 3 | GSE87822 | SRR4413794 | SRR4413795 | SRR4413796 | SRR4413797 |
| ChIP | ZHBTc4 | - | UNT | - | BRG1 | 1 | GSE87822 | SRR4413872 | SRR4413873 |  |  |
| ChIP | ZHBTc4 | - | DOX | - | BRG1 | 1 | GSE87822 | SRR4413878 | SRR4413879 |  |  |
| ChIP | ZHBTc4 | - | UNT | - | BRG1 | 2 | GSE87822 | SRR4413874 | SRR4413875 |  |  |
| ChIP | ZHBTc4 | - | DOX | - | BRG1 | 2 | GSE87822 | SRR4413880 | SRR4413881 |  |  |
| ChIP | ZHBTc4 | - | UNT | - | BRG1 | 3 | GSE87822 | SRR4413876 | SRR4413877 |  |  |
| ChIP | ZHBTc4 | - | DOX | - | BRG1 | 3 | GSE87822 | SRR4413882 | SRR4413883 |  |  |
| ChIP | ZHBTc4 | - | UNT | - | SOX2 | 1 | GSE87822 | SRR4413848 | SRR4413849 |  |  |
| ChIP | ZHBTc4 | - | DOX | - | SOX2 | 1 | GSE87822 | SRR4413854 | SRR4413855 |  |  |
| ChIP | ZHBTc4 | - | UNT | - | SOX2 | 2 | GSE87822 | SRR4413850 | SRR4413851 |  |  |
| ChIP | ZHBTc4 | - | DOX | - | SOX2 | 2 | GSE87822 | SRR4413856 | SRR4413857 |  |  |
| ChIP | ZHBTc4 | - | UNT | - | SOX2 | 3 | GSE87822 | SRR4413852 | SRR4413853 |  |  |
| ChIP | ZHBTc4 | - | DOX | - | SOX2 | 3 | GSE87822 | SRR4413859 | SRR4413859 |  |  |
| ChIP | ZHBTc4 | - | UNT | - | OCT4 | 1 | GSE87822 | SRR4413836 | SRR4413837 |  |  |
| ChIP | ZHBTc4 | - | DOX | - | OCT4 | 1 | GSE87822 | SRR4413842 | SRR4413843 |  |  |
| ChIP | ZHBTc4 | - | UNT | - | OCT4 | 2 | GSE87822 | SRR4413838 | SRR4413839 |  |  |
| ChIP | ZHBTc4 | - | DOX | - | OCT4 | 2 | GSE87822 | SRR4413844 | SRR4413845 |  |  |
| ChIP | ZHBTc4 | - | UNT | - | OCT4 | 3 | GSE87822 | SRR4413840 | SRR4413841 |  |  |
| ChIP | ZHBTc4 | - | DOX | - | OCT4 | 3 | GSE87822 | SRR4413846 | SRR4413847 |  |  |
| ChIP | ZHBTc4 | - | UNT | - | Input | 1 | GSE87822 | SRR4413884 | SRR4413885 |  |  |

**Table 5.** Description of publicly available pre-processed data used in this study.

| Accession | Sample number | Sample name | Factor | Cell_line | Source |
| --- | --- | --- | --- | --- | --- |
| GSE49848 | GSM1208217 | CME186_KLF4_CHIP-SEQ | KLF4 | ESC | cistromeDB |
| GSE57700 | GSM2065694 | SALL4-CHIP-IN-WT-OF-SALL-DKO | SALL4 | ESC | cistromeDB |
| GSE113915 | GSM3123484 | WT MESC NANOG | NANOG | ESC | cistromeDB |
| GSE90895 | GSM2417188 | ESC_ESRRB_CHIP-SEQ | ESRRB | ESC | cistromeDB |
| GSE99022 | GSM2630490 | WT_2I_H3K4ME3 | H3K4me3 | ESC | cistromeDB |
| GSE95781 | GSM2636047 | WT_H3K4ME1 | H3K4me1 | ESC | cistromeDB |
| GSE72886 | GSM1874094 | MESC_H3K27AC_CHIPSEQ | H3K27ac | ESC | cistromeDB |
| GSE111824 | GSM3398593 | WTTS_ATAc_rep1 | ATAC | TSC | GEO |
| GSE51511 | GSM1246722 | Sox2_ChIPseq_ZHBTc4-TSC | SOX2 | TSC | GEO |
| GSE74112 | GSM1910641 | PARP-1KO_ESC_Sox2_ChIPseq_Rep1 | SOX2 | ESC_PARP1KO | GEO |
| GSE74112 | GSM1910640 | WT_ESC_Sox2_ChIPseq_Rep1 | SOX2 | ESC | GEO |
| GSE87819 |  | GSE87819_mESC_BRG1fl_ATAc_72hrTAM_MERGED.DANPOS.bgsub.smooth | ATAC | ESC_BRG1fl | GEO |
| GSE87819 |  | GSE87819_mESC_BRG1fl_ATAc_UNT_MERGED.DANPOS.bgsub.smooth | ATAC | ESC_BRG1fl | GEO |
| GSE87821 |  | GSE87821_ZHBTc4.nucRNAseq.DESeq2_Results | RNA-seq | ZHBTc4 | GEO |
| GSE89599 |  | GSE89599_peaks_Asynchronous_ChIP_vs_Input_3rep_qval001_notInBlacklist | SOX2 | E14 | GEO |

### K-means clustering

Clusters in Fig. 3D were generated using the R function pheatmap with settings ‘clustering_distance_rows = “euclidean”, kmeans_k = 4’ on a matrix containing the log2 fold-change values in accessibility between MD-OCT4 and MD*-OCT4 at each cell cycle phase (columns) at each OCT4-bound locus (rows). Clusters were ordered according to the lowest mean log2 fold-change in EG1.

### Exponential curve fitting

Exponential decays were fitted using the R function nls with the formula y ∼ a*e^(-b*x)+c where a, b, and c are constants. Half-life values were derived as log(2)/b.

### Supplementary figure and table legends

**Supplementary Figure 1.**
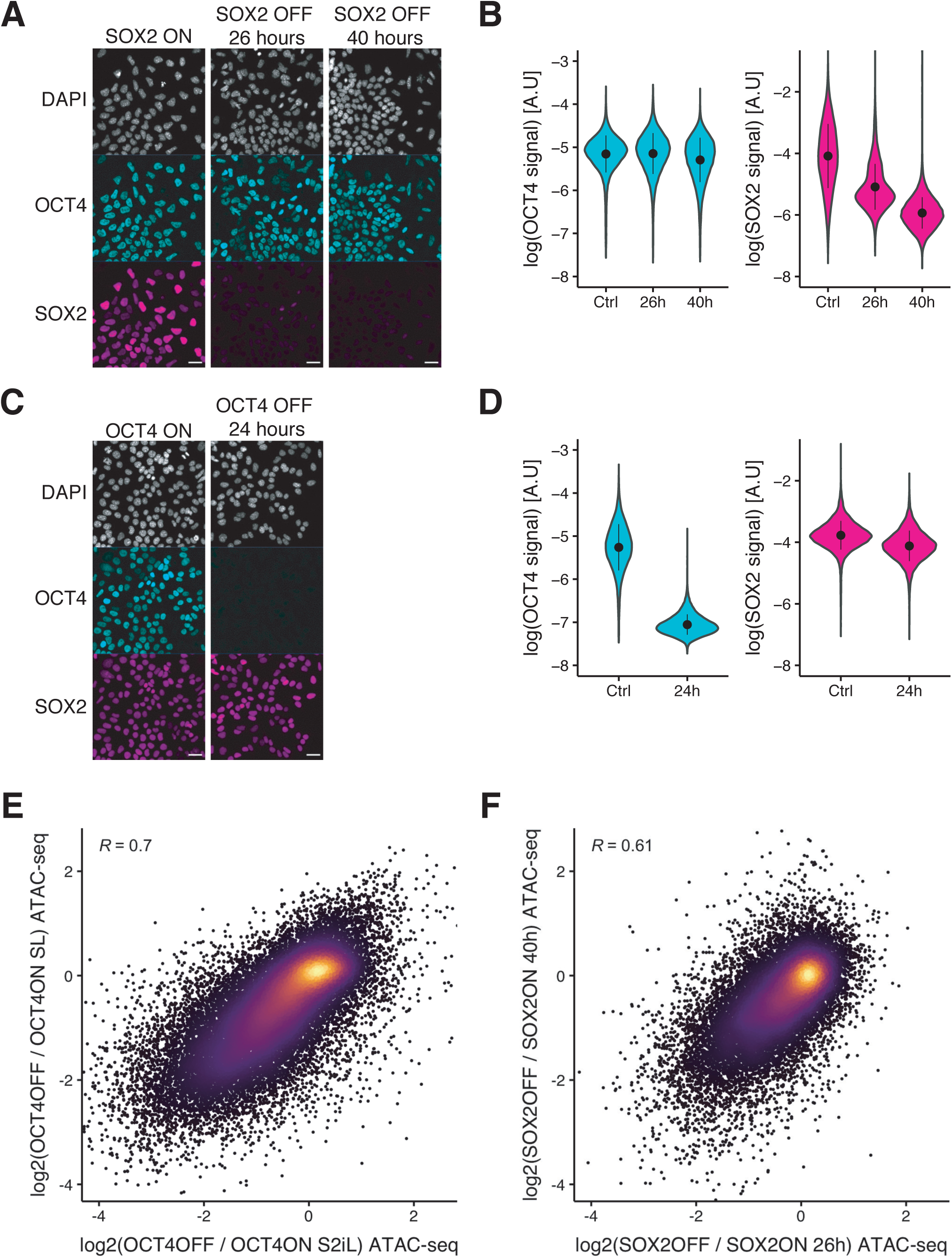
(A) Immunofluorescence of 2TS22C cells stained for DNA (DAPI), OCT4, and SOX2 without dox treatment (left), and after 26 hours (middle), and 40 hours (right) of dox treatment. (B) Violin plot of background-subtracted log values of immunofluorescence signal in OCT4 (left) and SOX2 (right) channels upon SOX2 depletion. Control: n=45’601 cells from 4 biological replicates including 2 technical replicates; 26 hours: n=42’298 cells from 3 biological replicates including 2 technical replicates; 40 hours: n= 32’342 cells from 2 technical replicates. Dots: mean; Vertical lines: standard deviation. (C) Immunofluorescence of ZHBTc4 cells stained for DNA (DAPI), OCT4, and SOX2 without dox treatment (left), and after 24 hours of dox treatment (right). (D) Violin plot of background-subtracted log values of immunofluorescence signal in OCT4 (left) and SOX2 (right) channels upon OCT4 depletion. Control: n=26’119 cells from 3 biological replicates. 24 hours: n=23’157 cells from 3 biological replicates. Dots: mean; Vertical lines: standard deviation. (E) Correlation between the log2 fold-change values of accessibility upon OCT4 depletion in S2iL (x-axis) and SL (y-axis) at OCT4-bound sites. (F) Correlation between the log2 fold-change values of accessibility upon SOX2 depletion after 26 hours (x-axis) and 40 hours (y-axis) of dox treatment at SOX2 binding sites. R is Pearson correlation coefficient. Scale bars: 30 μm.

**Supplementary Figure 2.**
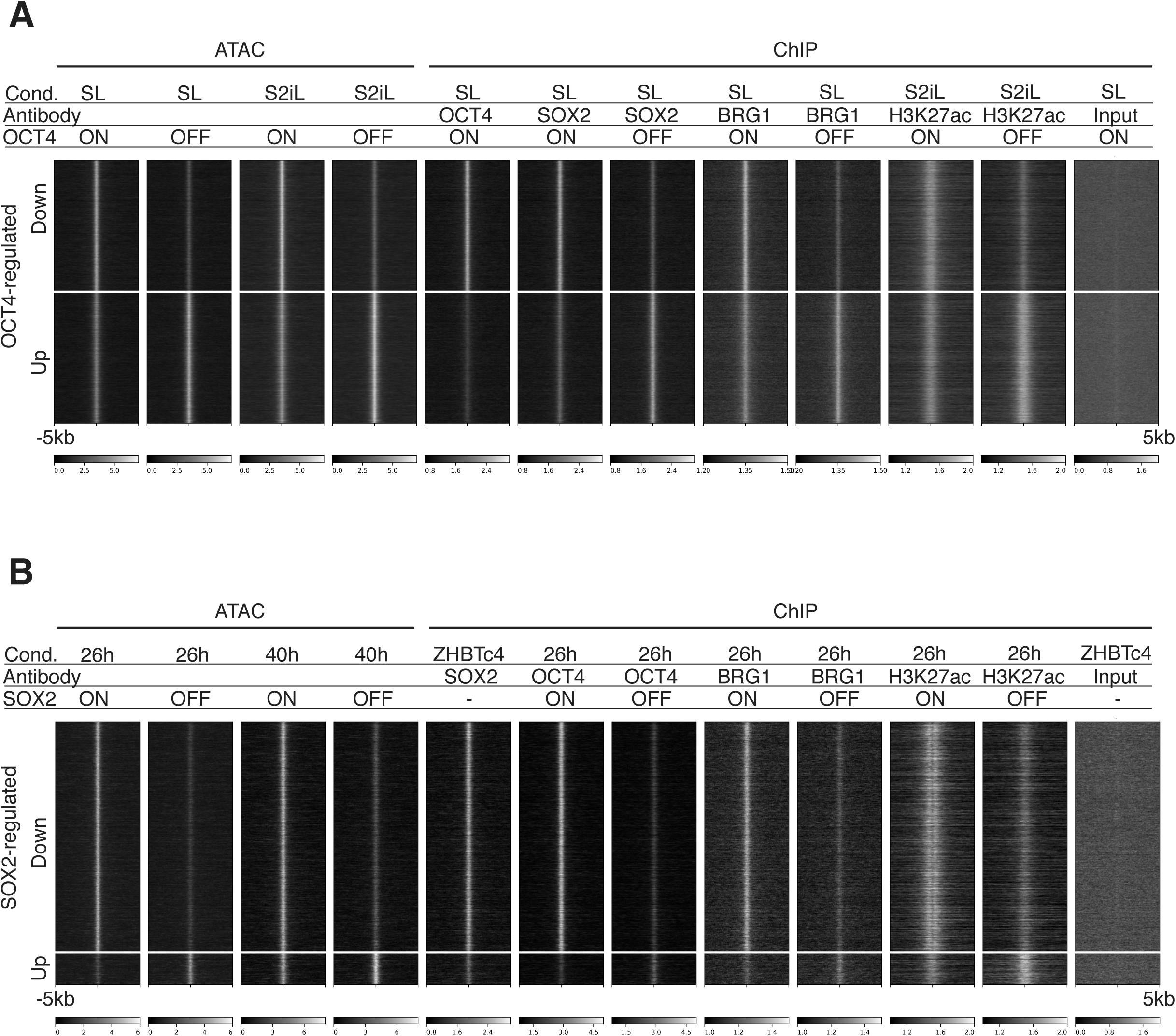
Heatmaps of RPKM-normalized ATAC-seq and ChIP-seq binding profiles upon OCT4 (A) and SOX2 (B) depletion 5 kb around OCT4-regulated (A) and SOX2-regulated (B) loci. Each row represents one individual locus and each column represents one experimental condition.

**Supplementary Figure 3.**
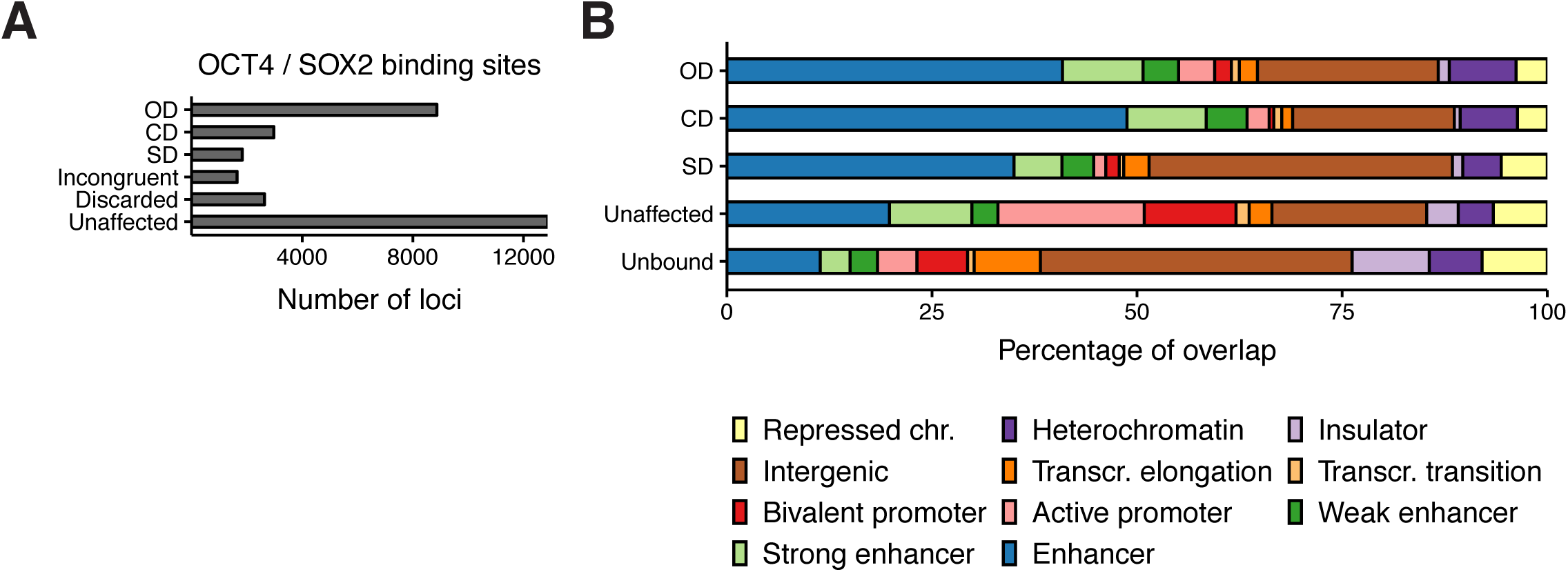
(A) Classification of all OCT4 and SOX2 binding sites into OD, CD, and SD loci as well as loci that were discarded due to differences in untreated cells between conditions or cell lines (Discarded), due to incongruent effect on accessibility after depletion in different conditions (Incongruent) and those that were unaffected by depletion. (B) ChromHMM signal enrichment at OD, CD, and SD loci as well as loci that were unaffected by depletion or not bound by OCT4 or SOX2.

**Supplementary Figure 4.**
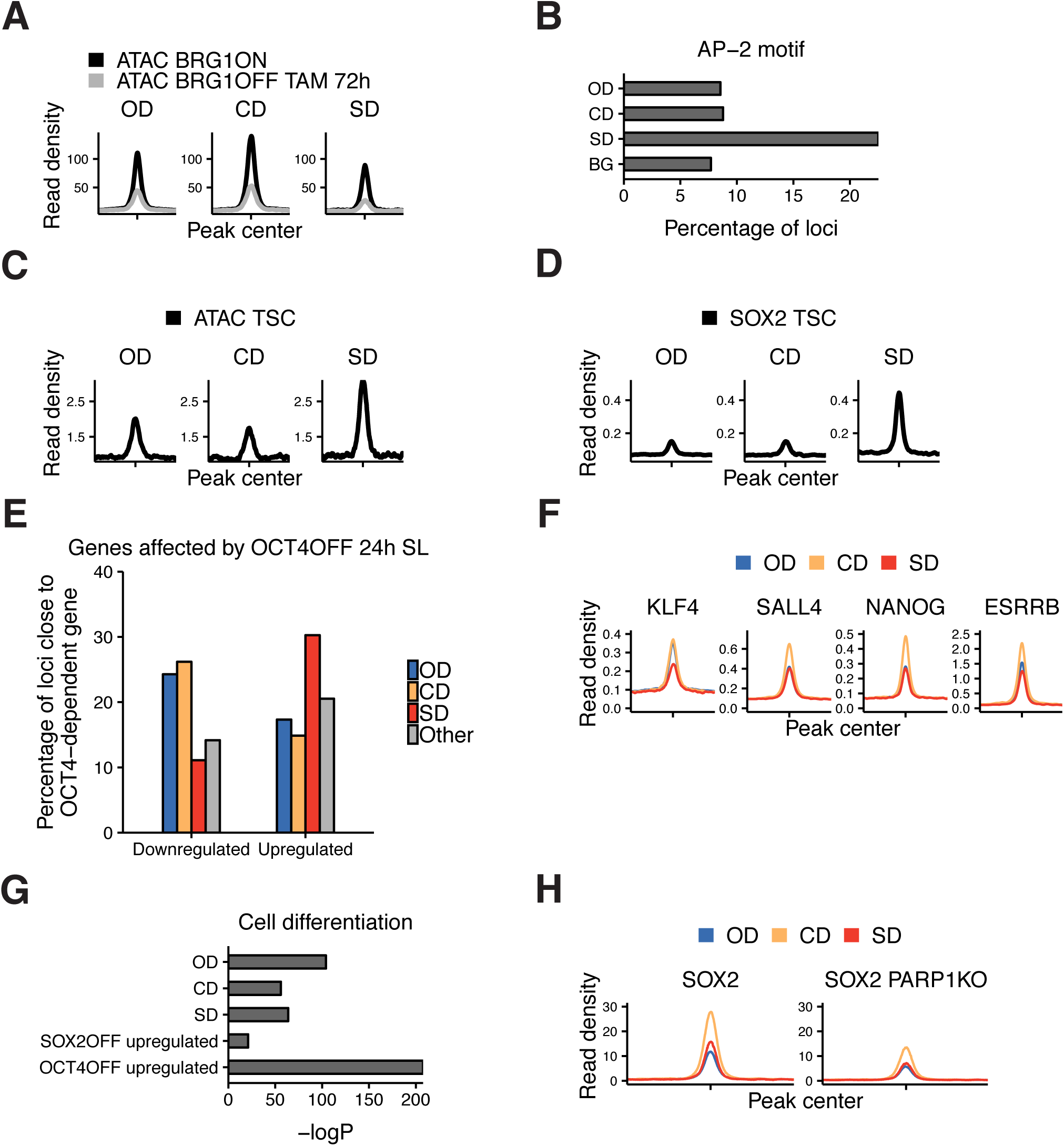
(A) Average ATAC-seq signal 2 kb around OD, CD, and SD loci in BRG1fl cells that were treated with tamoxifen (TAM) or left untreated. (B) Frequency of the AP-2 motif 2 kb around OD, CD, and SD loci, and in background regions (BG). (C) Average SOX2 ChIP-seq signal in TS cells 2 kb around OD, CD, and SD loci. (D) Average ATAC-seq signal in TS cells 2 kb around OD, CD, and SD loci. (E) Percentage of the closest gene in the OD, CD, and SD groups as well as all other accessible regions (Other) whose nascent RNA levels are downregulated or upregulated upon 24 hours of OCT4 depletion (F) Average ChIP-seq signal of ESRRB, NANOG, KLF4, and SALL4 in ES cells 2 kb around OD, CD, and SD loci. (G) Enrichment (-log(p)) values for the closest gene in the OD, CD, and SD groups in the “Cell differentiation” gene ontology set. (H) SOX2 binding profiles 2 kb around OD, CD, and SD loci in wt and PARP1 KO ES cells.

**Supplementary Figure 5.**
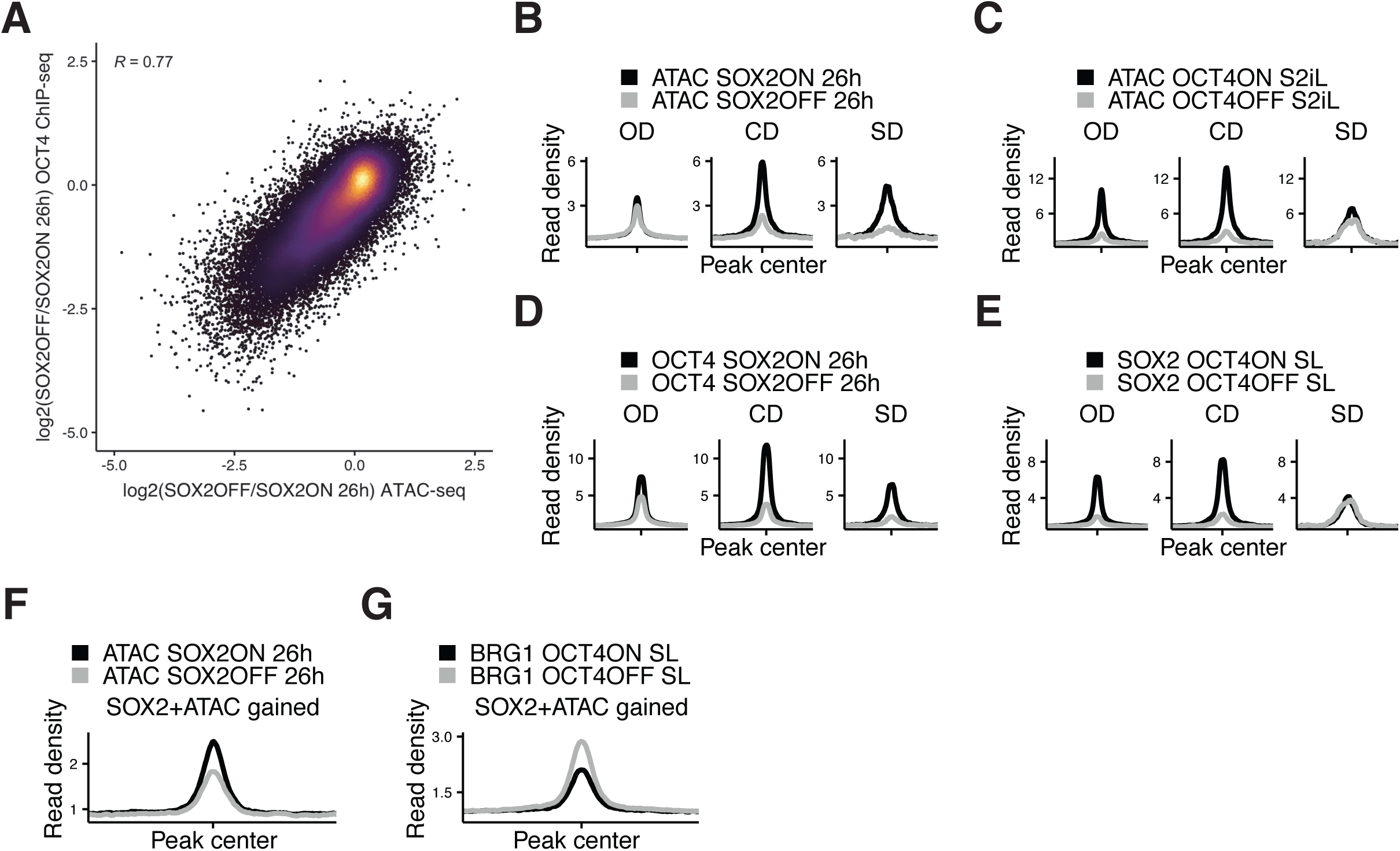
(A) Correlation between log2 fold-change values of accessibility (x-axis) and OCT4 binding (y-axis) upon SOX2 depletion in 2TS22C cells with dox treatment for 26 hours. R is Pearson correlation coefficient. (B-C) Average RPKM-normalized ATAC-seq signal 2 kb around OD (n=3’730), CD (n=1’463), and SD (n=273) loci that overlap with a canonical OCT4::SOX2 motif upon SOX2 (B) and OCT4 (C) depletion. (D-E) Average RPKM-normalized OCT4 (D) and SOX2 (E) ChIP-seq signal 2 kb around OD, CD, and SD loci that overlap with a canonical OCT4::SOX2 motif upon SOX2 (D) and OCT4 (E) depletion. (F) Average RPKM-normalized ATAC-seq signal upon SOX2 depletion 2 kb around loci that display a significant increase in accessibility and SOX2 binding upon OCT4 depletion. (G) Average RPKM-normalized BRG1 ChIP-seq signal upon OCT4 depletion 2 kb around loci that display a significant increase in accessibility and SOX2 binding upon OCT4 depletion.

**Supplementary Figure 6.**
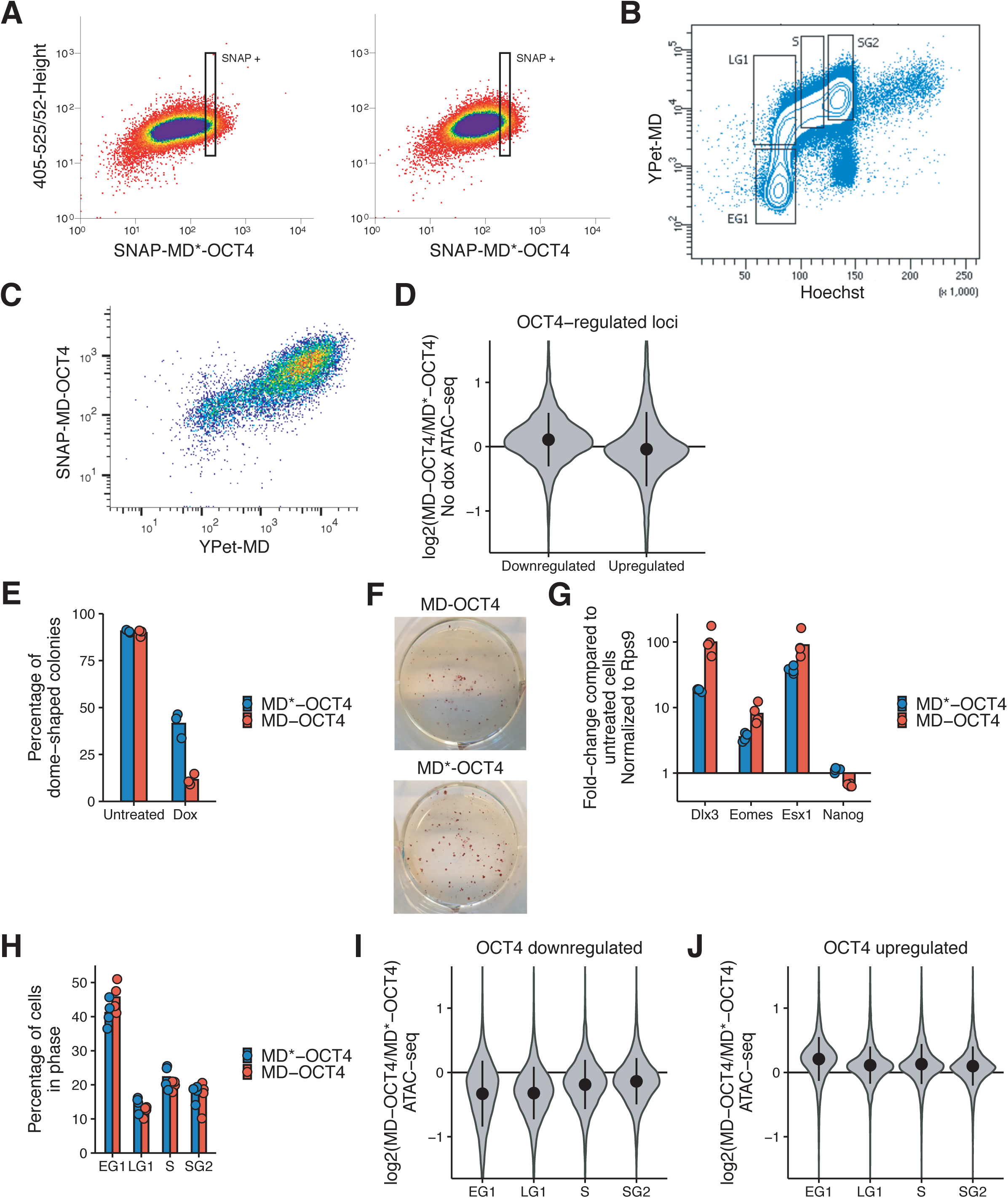
(A) Gate used to sort SNAP-MD-OCT4 (left) and SNAP-MD*-OCT4 (right) cells for the same average SNAP-Cell 647-SiR signal. Y-axis: Signal amplitude at 405 nm excitation and 526/52 nm emission (negative control). X-axis: Signal amplitude at 640 nm excitation and 671/30 nm emission (SNAP signal). (B) Example of a sorting experiment for different phases of the cell cycle in cells expressing YPet-MD and stained for Hoechst33258. Y-axis: Integrated signal at 488 nm excitation and 525/50 nm emission (YPet). X-axis: Signal amplitude at 355 nm excitation and 450/50 nm emission (Hoechst) (C) Correlation between YPet-MD and SNAP-MD-OCT4 expression in MD-OCT4 cells as measured by flow cytometry. Y-axis: Integrated signal at 640 nm excitation and 670/14 nm emission (SNAP). X-axis: Integrated signal at 488 nm excitation and 525/50 nm emission (YPet). (D) Violin plot of log2 fold-change values of accessibility between MD-OCT4 and MD*-OCT4 cells in significantly downregulated and upregulated loci (see Fig.1B) in unsorted cells in the absence of dox. Dots: mean; Vertical lines: standard deviation. (E) Percentage of dome-shaped colonies as assessed by microscopy in the ZHBTc4 cell line upon dox treatment and with overexpression of SNAP-MD*-OCT4 or SNAP-MD-OCT4. n=3 biological replicates. (F) Representative alkaline phosphatase staining from cells in (E). (G) Fold-change of expression levels of differentiation markers (Dlx3, Eomes and Esx1) and Nanog, measured by qRT-PCR in dox-treated versus untreated cells, in MD-OCT4 and MD*-OCT4 cells. Each sample is normalized to the expression of Rps9. n=4 biological replicates. (H) Percentage of cells in EG1/LG1/S/SG2 phases as determined by flow cytometry in MD-OCT4 and MD*-OCT4 cells. n=4 biological replicates. (I-J) Violin plot of log2 fold-change values of accessibility between MD-OCT4 and MD*-OCT4 cells in different cell cycle phases at significantly downregulated (I) and upregulated (J) loci (see Fig. 1B). Dots: mean; Vertical lines: standard deviation.

**Supplementary Figure 7.**
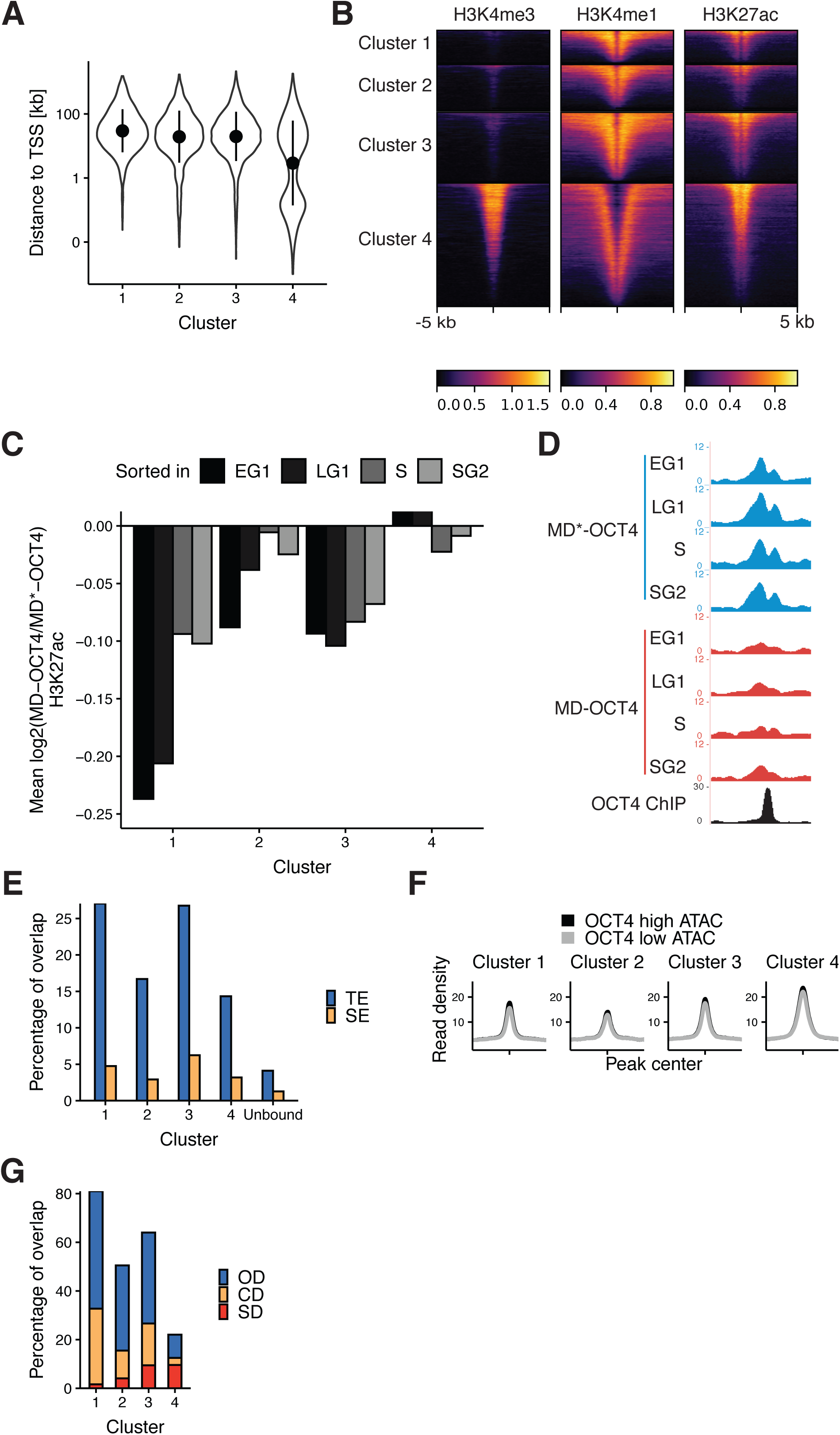
(A) Violin plot of distance to closest TSS in clusters from Fig. 3D. Dots: mean; Vertical lines: standard deviation. (B) Heatmap of ChIP-seq signal of H3K4me3, H3K4me1, and H3K27ac in wt ES cells in the different clusters. (C) Average log2 fold-change values of H3K27ac ChIP-seq signal between MD-OCT4 and MD*-OCT4 cells in the different clusters (including 500bp flanking regions at each side) at different cell cycle stages. (D) Genome browser tracks of RPKM-normalized H3K27ac profiles across the cell cycle for a cluster 1 locus (chr11:6894809-6895533) that decreases in accessibility and H3K27ac upon transient OCT4 depletion in M-G1. (E) Percentage of loci in the different clusters and at non-OCT4 bound regions overlapping typical enhancers (TE) and super-enhancers (SE) in mouse ES cells. (F) ATAC-seq signal in cells sorted for high and low endogenous OCT4 levels in the different clusters. (G) Percentage of loci in the different clusters overlapping OD, CD, and SD loci (see Fig. 1C).

**Supplementary Figure 8.**
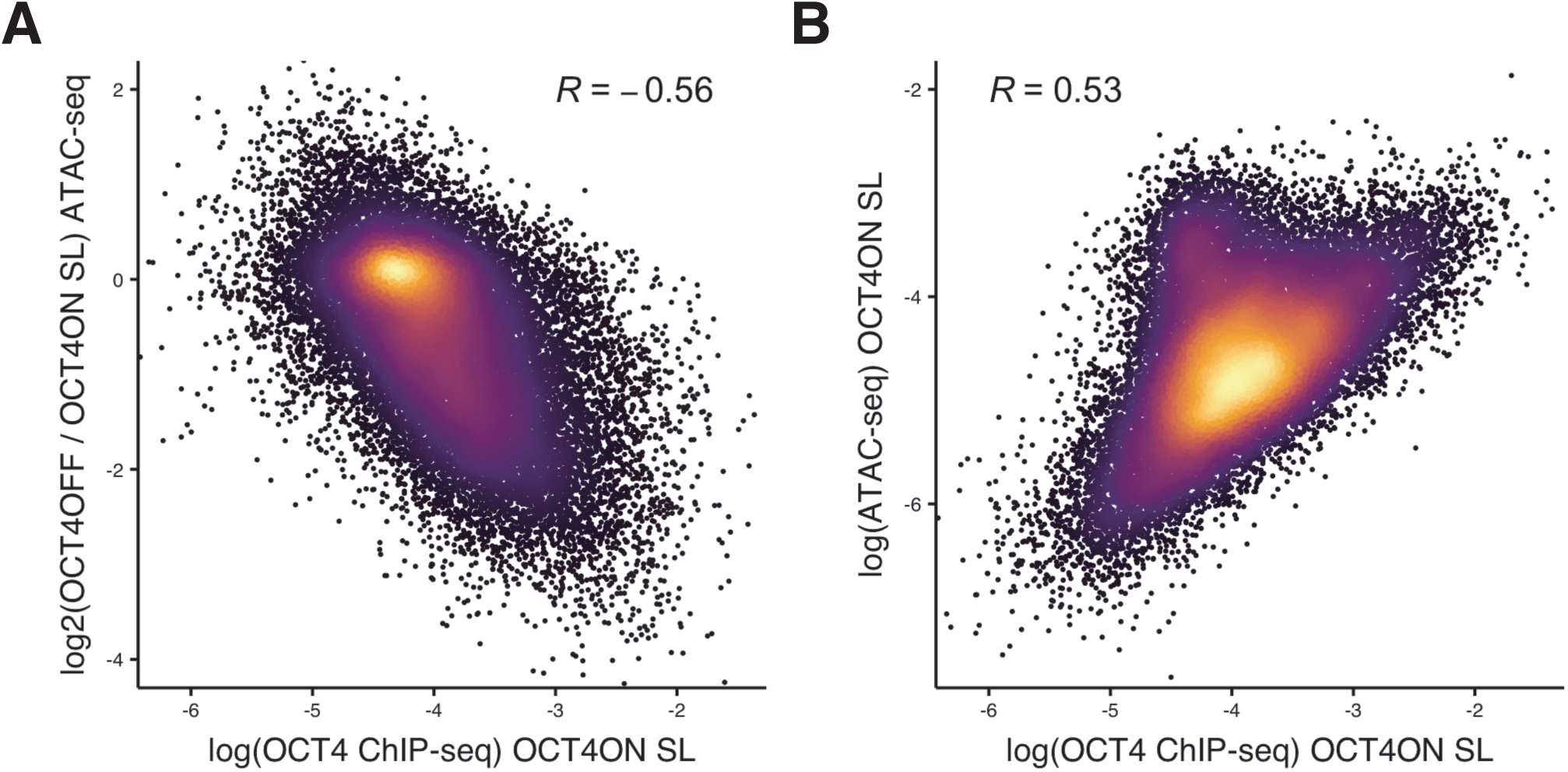
(A) Correlation between the log of normalized OCT4 ChIP-seq reads per bp (x-axis) and the log2 fold-change values of accessibility loss upon OCT4 depletion (y-axis) at all OCT4 binding sites in ZHBTc4 cells. (B) Correlation between the log of normalized OCT4 ChIP-seq reads per bp (x-axis) and the log of normalized ATAC-seq reads per bp (y-axis) at all OCT4 binding sites in ZHBTc4 cells. R is Pearson correlation coefficient.

**Supplementary Figure 9.**
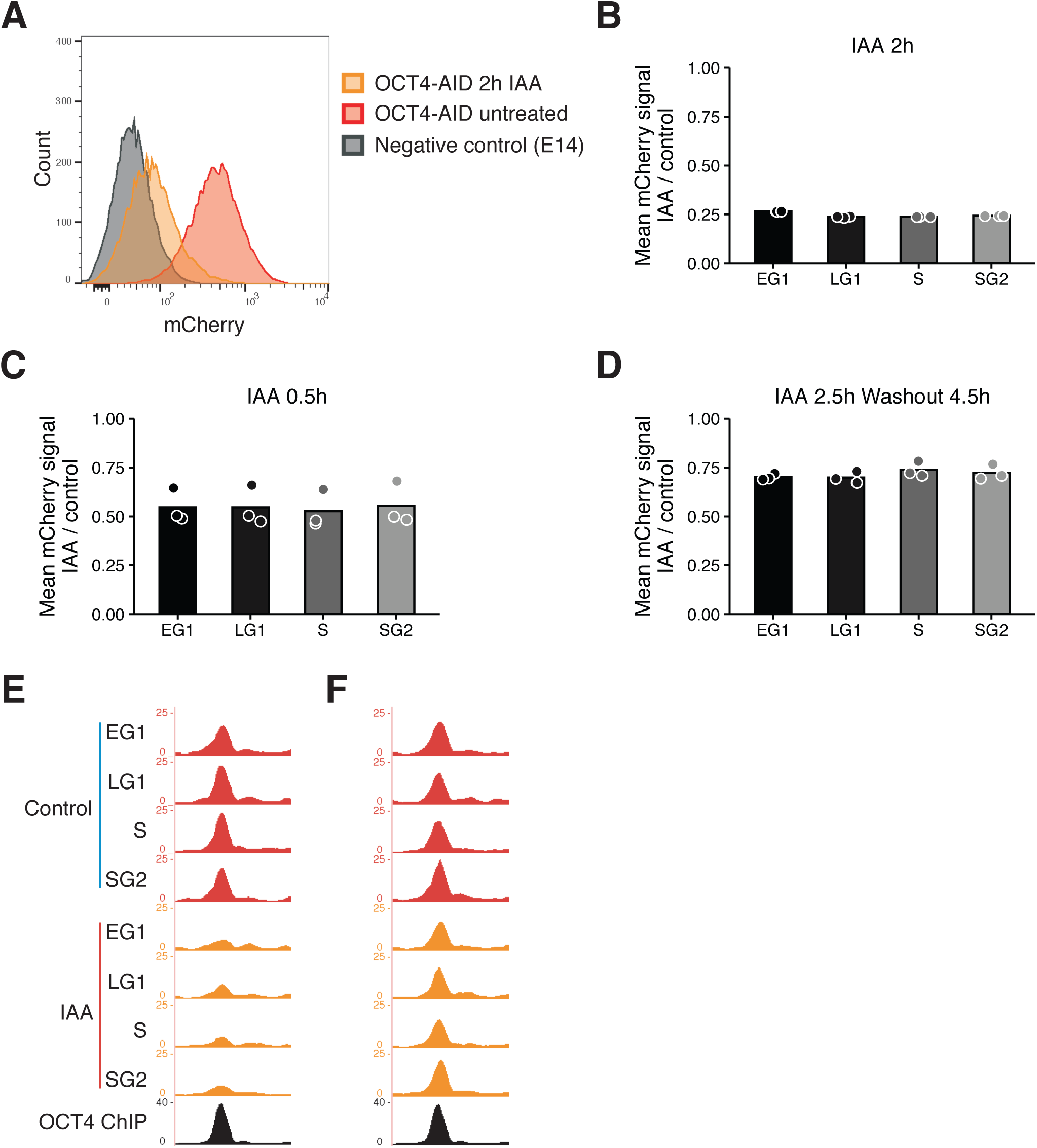
(A) Histogram of mCherry signal in untreated mCherry OCT4-AID cells and treated with IAA for 2 hours as well as mCherry negative E14 ES cells as measured by flow cytometry. X-axis: Integrated signal at 561 nm excitation and 610/20 nm emission. Y-axis: Counts. (B-D) Fold-change of red fluorescence (mCherry) signal between treated and untreated mCherry-OCT4-AID cells as determined by flow cytometry in different cell cycle phases upon 2 h IAA treatment (B), 0.5 h IAA treatment (C), and after 2.5 h IAA treatment followed by 4.5 h of washout (D). n=3 biological replicates. (E-F) Genome browser tracks of accessibility profiles of a cluster 1 locus at chr10:95455826-95456819 after IAA treatment (E) and after IAA treatment followed by washout (F).

## References

1. Adachi K, Nikaido I, Ohta H, Ohtsuka S, Ura H, Kadota M, Wakayama T, Ueda HR, Niwa H. 2013. Context-Dependent Wiring of Sox2 Regulatory Networks for Self-Renewal of Embryonic and Trophoblast Stem Cells. Molecular Cell 52:380–392. doi:10.1016/j.molcel.2013.09.002

2. Aksoy I, Giudice V, Delahaye E, Wianny F, Aubry M, Mure M, Chen J, Jauch R, Bogu GK, Nolden T, Himmelbauer H, Xavier Doss M, Sachinidis A, Schulz H, Hummel O, Martinelli P, Hübner N, Stanton LW, Real FX, Bourillot P-Y, Savatier P. 2014. Klf4 and Klf5 differentially inhibit mesoderm and endoderm differentiation in embryonic stem cells. Nat Commun 5:3719. doi:10.1038/ncomms4719

3. Alber AB, Paquet ER, Biserni M, Naef F, Suter DM. 2018. Single Live Cell Monitoring of Protein Turnover Reveals Intercellular Variability and Cell-Cycle Dependence of Degradation Rates. Mol Cell 71:1079–1091.e9. doi:10.1016/j.molcel.2018.07.023

4. Boyer LA, Lee TI, Cole MF, Johnstone SE, Levine SS, Zucker JP, Guenther MG, Kumar RM, Murray HL, Jenner RG, Gifford DK, Melton DA, Jaenisch R, Young RA. 2005. Core Transcriptional Regulatory Circuitry in Human Embryonic Stem Cells. Cell 122:947–956. doi:10.1016/j.cell.2005.08.020

5. Buenrostro JD, Giresi PG, Zaba LC, Chang HY, Greenleaf WJ. 2013. Transposition of native chromatin for fast and sensitive epigenomic profiling of open chromatin, DNA-binding proteins and nucleosome position. Nat Methods 10:1213–1218. doi:10.1038/nmeth.2688

6. Cao K, Collings CK, Morgan MA, Marshall SA, Rendleman EJ, Ozark PA, Smith ER, Shilatifard A. 2018. An Mll4/COMPASS-Lsd1 epigenetic axis governs enhancer function and pluripotency transition in embryonic stem cells. Sci Adv 4:eaap8747. doi:10.1126/sciadv.aap8747

7. Carpenter AE, Jones TR, Lamprecht MR, Clarke C, Kang IH, Friman O, Guertin DA, Chang JH, Lindquist RA, Moffat J, Golland P, Sabatini DM. 2006. CellProfiler: image analysis software for identifying and quantifying cell phenotypes. Genome Biol 7:R100. doi:10.1186/gb-2006-7-10-r100

8. Chen J, Zhang Z, Li L, Chen B-C, Revyakin A, Hajj B, Legant W, Dahan M, Lionnet T, Betzig E, Tjian R, Liu Z. 2014. Single-Molecule Dynamics of Enhanceosome Assembly in Embryonic Stem Cells. Cell 156:1274–1285. doi:10.1016/j.cell.2014.01.062

9. Chen W, Gardeux V, Meireles-Filho A, Deplancke B. 2017. Profiling of Single-Cell Transcriptomes. Curr Protoc Mouse Biol 7:145–175. doi:10.1002/cpmo.30

10. Chronis C, Fiziev P, Papp B, Butz S, Bonora G, Sabri S, Ernst J, Plath K. 2017. Cooperative Binding of Transcription Factors Orchestrates Reprogramming. Cell 168:442–459.e20. doi:10.1016/j.cell.2016.12.016

11. Cirillo LA, Lin FR, Cuesta I, Friedman D, Jarnik M, Zaret KS. 2002. Opening of Compacted Chromatin by Early Developmental Transcription Factors HNF3 (FoxA) and GATA-4. Molecular Cell 9:279–289. doi:10.1016/S1097-2765(02)00459-8

12. Creyghton MP, Cheng AW, Welstead GG, Kooistra T, Carey BW, Steine EJ, Hanna J, Lodato MA, Frampton GM, Sharp PA, Boyer LA, Young RA, Jaenisch R. 2010. Histone H3K27ac separates active from poised enhancers and predicts developmental state. Proc Natl Acad Sci U S A 107:21931–21936. doi:10.1073/pnas.1016071107

13. Deluz C, Friman ET, Strebinger D, Benke A, Raccaud M, Callegari A, Leleu M, Manley S, Suter DM. 2016. A role for mitotic bookmarking of SOX2 in pluripotency and differentiation. Genes Dev 30:2538–2550. doi:10.1101/gad.289256.116

14. Dharmasiri N, Dharmasiri S, Estelle M. 2005. The F-box protein TIR1 is an auxin receptor. Nature 435:441. doi:10.1038/nature03543

15. Dobin A, Davis CA, Schlesinger F, Drenkow J, Zaleski C, Jha S, Batut P, Chaisson M, Gingeras TR. 2013. STAR: ultrafast universal RNA-seq aligner. Bioinformatics 29:15–21. doi:10.1093/bioinformatics/bts635

16. Dull T, Zufferey R, Kelly M, Mandel RJ, Nguyen M, Trono D, Naldini L. 1998. A Third-Generation Lentivirus Vector with a Conditional Packaging System. Journal of Virology 72:8463–8471.

17. Dunn SJ, Martello G, Yordanov B, Emmott S, Smith AG. 2014. Defining an essential transcription factor program for naive pluripotency. Science 344:1156–60. doi:10.1126/science.1248882

18. Durinck S, Moreau Y, Kasprzyk A, Davis S, De Moor B, Brazma A, Huber W. 2005. BioMart and Bioconductor: a powerful link between biological databases and microarray data analysis. Bioinformatics 21:3439–3440. doi:10.1093/bioinformatics/bti525

19. Edgar R, Domrachev M, Lash AE. 2002. Gene Expression Omnibus: NCBI gene expression and hybridization array data repository. Nucleic Acids Res 30:207–210. doi:10.1093/nar/30.1.207

20. Egli D, Birkhoff G, Eggan K. 2008. Mediators of reprogramming: transcription factors and transitions through mitosis. Nat Rev Mol Cell Biol 9:505–516. doi:10.1038/nrm2439

21. ENCODE Project Consortium. 2012. An integrated encyclopedia of DNA elements in the human genome. Nature 489:57–74. doi:10.1038/nature11247

22. Festuccia N, Owens N, Papadopoulou T, Gonzalez I, Tachtsidi A, Vandoermel-Pournin S, Gallego E, Gutierrez N, Dubois A, Cohen-Tannoudji M, Navarro P. 2019. Transcription factor activity and nucleosome organization in mitosis. Genome Res 29:250–260. doi:10.1101/gr.243048.118

23. Gao F, Kwon SW, Zhao Y, Jin Y. 2009. PARP1 Poly(ADP-ribosyl)ates Sox2 to Control Sox2 Protein Levels and FGF4 Expression during Embryonic Stem Cell Differentiation. J Biol Chem 284:22263–22273. doi:10.1074/jbc.M109.033118

24. Goldstein I, Paakinaho V, Baek S, Sung M-H, Hager GL. 2017. Synergistic gene expression during the acute phase response is characterized by transcription factor assisted loading. Nat Commun 8:1849. doi:10.1038/s41467-017-02055-5

25. Gu Z, Eils R, Schlesner M. 2016. Complex heatmaps reveal patterns and correlations in multidimensional genomic data. Bioinformatics 32:2847–2849. doi:10.1093/bioinformatics/btw313

26. Heinz S, Benner C, Spann N, Bertolino E, Lin YC, Laslo P, Cheng JX, Murre C, Singh H, Glass CK. 2010. Simple combinations of lineage-determining transcription factors prime cis-regulatory elements required for macrophage and B cell identities. Mol Cell 38:576–589. doi:10.1016/j.molcel.2010.05.004

27. Hinrichs AS, Karolchik D, Baertsch R, Barber GP, Bejerano G, Clawson H, Diekhans M, Furey TS, Harte RA, Hsu F, Hillman-Jackson J, Kuhn RM, Pedersen JS, Pohl A, Raney BJ, Rosenbloom KR, Siepel A, Smith KE, Sugnet CW, Sultan-Qurraie A, Thomas DJ, Trumbower H, Weber RJ, Weirauch M, Zweig AS, Haussler D, Kent WJ. 2006. The UCSC Genome Browser Database: update 2006. Nucleic Acids Res 34:D590–598. doi:10.1093/nar/gkj144

28. Ho L, Miller EL, Ronan JL, Ho WQ, Jothi R, Crabtree GR. 2011. esBAF facilitates pluripotency by conditioning the genome for LIF/STAT3 signalling and by regulating polycomb function. Nature Cell Biology 13:903–913. doi:10.1038/ncb2285

29. Holland AJ, Fachinetti D, Han JS, Cleveland DW. 2012. Inducible, reversible system for the rapid and complete degradation of proteins in mammalian cells. PNAS 109:E3350–E3357. doi:10.1073/pnas.1216880109

30. Hsiung CC-S, Morrissey CS, Udugama M, Frank CL, Keller CA, Baek S, Giardine B, Crawford GE, Sung M-H, Hardison RC, Blobel GA. 2015. Genome accessibility is widely preserved and locally modulated during mitosis. Genome Res 25:213–225. doi:10.1101/gr.180646.114

31. Ishiuchi T, Ohishi H, Sato T, Kamimura S, Yorino M, Abe S, Suzuki A, Wakayama T, Suyama M, Sasaki H. 2019. Zfp281 Shapes the Transcriptome of Trophoblast Stem Cells and Is Essential for Placental Development. Cell Rep 27:1742–1754.e6. doi:10.1016/j.celrep.2019.04.028

32. Iwafuchi-Doi M, Zaret KS. 2014. Pioneer transcription factors in cell reprogramming. Genes Dev 28:2679–2692. doi:10.1101/gad.253443.114

33. Jacobs J, Atkins M, Davie K, Imrichova H, Romanelli L, Christiaens V, Hulselmans G, Potier D, Wouters J, Taskiran II, Paciello G, González-Blas CB, Koldere D, Aibar S, Halder G, Aerts S. 2018. The transcription factor Grainy head primes epithelial enhancers for spatiotemporal activation by displacing nucleosomes. Nat Genet 50:1011–1020. doi:10.1038/s41588-018-0140-x

34. Kadauke S, Udugama MI, Pawlicki JM, Achtman JC, Jain DP, Cheng Y, Hardison RC, Blobel GA. 2012. Tissue-specific mitotic bookmarking by hematopoietic transcription factor GATA1. Cell 150:725–737. doi:10.1016/j.cell.2012.06.038

35. Karwacki-Neisius V, Göke J, Osorno R, Halbritter F, Ng JH, Weiße AY, Wong FCK, Gagliardi A, Mullin NP, Festuccia N, Colby D, Tomlinson SR, Ng H-H, Chambers I. 2013. Reduced Oct4 Expression Directs a Robust Pluripotent State with Distinct Signaling Activity and Increased Enhancer Occupancy by Oct4 and Nanog. Cell Stem Cell 12:531–545. doi:10.1016/j.stem.2013.04.023

36. Kent WJ, Sugnet CW, Furey TS, Roskin KM, Pringle TH, Zahler AM, Haussler D. 2002. The human genome browser at UCSC. Genome Res 12:996–1006. doi:10.1101/gr.229102

37. Kepinski S, Leyser O. 2005. The Arabidopsis F-box protein TIR1 is an auxin receptor. Nature 435:446–451. doi:10.1038/nature03542

38. Kim K-Y, Tanaka Y, Su J, Cakir B, Xiang Y, Patterson B, Ding J, Jung Y-W, Kim J-H, Hysolli E, Lee H, Dajani R, Kim J, Zhong M, Lee J-H, Skalnik D, Lim JM, Sullivan GJ, Wang J, Park I-H. 2018. Uhrf1 regulates active transcriptional marks at bivalent domains in pluripotent stem cells through Setd1a. Nat Commun 9:2583. doi:10.1038/s41467-018-04818-0

39. King HW, Klose RJ. 2017. The pioneer factor OCT4 requires the chromatin remodeller BRG1 to support gene regulatory element function in mouse embryonic stem cells. Elife 6. doi:10.7554/eLife.22631

40. Kumar V, Rayan NA, Muratani M, Lim S, Elanggovan B, Xin L, Lu T, Makhija H, Poschmann J, Lufkin T, Ng HH, Prabhakar S. 2016. Comprehensive benchmarking reveals H2BK20 acetylation as a distinctive signature of cell-state-specific enhancers and promoters. Genome Res 26:612–623. doi:10.1101/gr.201038.115

41. Lawrence M, Huber W, Pagès H, Aboyoun P, Carlson M, Gentleman R, Morgan MT, Carey VJ. 2013. Software for computing and annotating genomic ranges. PLoS Comput Biol 9:e1003118. doi:10.1371/journal.pcbi.1003118

42. Lee J, Go Y, Kang I, Han Y-M, Kim J. 2010. Oct-4 controls cell-cycle progression of embryonic stem cells. Biochem J 426:171–181. doi:10.1042/BJ20091439

43. Li H, Handsaker B, Wysoker A, Fennell T, Ruan J, Homer N, Marth G, Abecasis G, Durbin R. 2009. The Sequence Alignment/Map format and SAMtools. Bioinformatics 25:2078– 2079. doi:10.1093/bioinformatics/btp352

44. Li S, Zheng EB, Zhao L, Liu S. 2019. Nonreciprocal and Conditional Cooperativity Directs the Pioneer Activity of Pluripotency Transcription Factors. bioRxiv 633826. doi:10.1101/633826

45. Liu Y, Pelham-Webb B, Di Giammartino DC, Li J, Kim D, Kita K, Saiz N, Garg V, Doane A, Giannakakou P, Hadjantonakis A-K, Elemento O, Apostolou E. 2017. Widespread Mitotic Bookmarking by Histone Marks and Transcription Factors in Pluripotent Stem Cells. Cell Rep 19:1283–1293. doi:10.1016/j.celrep.2017.04.067

46. Liu Z, Kraus WL. 2017. Catalytic-Independent Functions of PARP-1 Determine Sox2 Pioneer Activity at Intractable Genomic Loci. Molecular Cell 65:589–603.e9. doi:10.1016/j.molcel.2017.01.017

47. Lukinavičius G, Umezawa K, Olivier N, Honigmann A, Yang G, Plass T, Mueller V, Reymond L, Corrêa IR, Luo Z-G, Schultz C, Lemke EA, Heppenstall P, Eggeling C, Manley S, Johnsson K. 2013. A near-infrared fluorophore for live-cell super-resolution microscopy of cellular proteins. Nat Chem 5:132–139. doi:10.1038/nchem.1546

48. Masui S, Nakatake Y, Toyooka Y, Shimosato D, Yagi R, Takahashi K, Okochi H, Okuda A, Matoba R, Sharov AA, Ko MS, Niwa H. 2007. Pluripotency governed by Sox2 via regulation of Oct3/4 expression in mouse embryonic stem cells. Nature cell biology 9:625–35. doi:10.1038/ncb1589

49. Mayran A, Khetchoumian K, Hariri F, Pastinen T, Gauthier Y, Balsalobre A, Drouin J. 2018. Pioneer factor Pax7 deploys a stable enhancer repertoire for specification of cell fate. Nat Genet 50:259–269. doi:10.1038/s41588-017-0035-2

50. Mayran A, Sochodolsky K, Khetchoumian K, Harris J, Gauthier Y, Bemmo A, Balsalobre A, Drouin J. 2019. Pioneer and nonpioneer cooperation drives lineage specific chromatin opening. bioRxiv 472647. doi:10.1101/472647

51. McDaniel SL, Gibson TJ, Schulz KN, Fernandez Garcia M, Nevil M, Jain SU, Lewis PW, Zaret KS, Harrison MM. 2019. Continued Activity of the Pioneer Factor Zelda Is Required to Drive Zygotic Genome Activation. Mol Cell 74:185–195.e4. doi:10.1016/j.molcel.2019.01.014

52. Mei S, Qin Q, Wu Q, Sun H, Zheng R, Zang C, Zhu M, Wu J, Shi X, Taing L, Liu T, Brown M, Meyer CA, Liu XS. 2017. Cistrome Data Browser: a data portal for ChIP-Seq and chromatin accessibility data in human and mouse. Nucleic Acids Res 45:D658–D662. doi:10.1093/nar/gkw983

53. Mistri TK, Arindrarto W, Ng WP, Wang C, Lim LH, Sun L, Chambers I, Wohland T, Robson P. 2018. Dynamic changes in Sox2 spatio-temporal expression promote the second cell fate decision through Fgf4/Fgfr2 signaling in preimplantation mouse embryos. Biochem J 475:1075–1089. doi:10.1042/BCJ20170418

54. Mistri TK, Devasia AG, Chu LT, Ng WP, Halbritter F, Colby D, Martynoga B, Tomlinson SR, Chambers I, Robson P, Wohland T. 2015. Selective influence of Sox2 on POU transcription factor binding in embryonic and neural stem cells. EMBO Rep 16:1177–1191. doi:10.15252/embr.201540467

55. Morawska M, Ulrich HD. 2013. An expanded tool kit for the auxin-inducible degron system in budding yeast. Yeast 30:341–351. doi:10.1002/yea.2967

56. Nabet B, Roberts JM, Buckley DL, Paulk J, Dastjerdi S, Yang A, Leggett AL, Erb MA, Lawlor MA, Souza A, Scott TG, Vittori S, Perry JA, Qi J, Winter GE, Wong K-K, Gray NS, Bradner JE. 2018. The dTAG system for immediate and target-specific protein degradation. Nat Chem Biol 14:431–441. doi:10.1038/s41589-018-0021-8

57. Nishimoto M, Fukushima A, Okuda A, Muramatsu M. 1999. The Gene for the Embryonic Stem Cell Coactivator UTF1 Carries a Regulatory Element Which Selectively Interacts with a Complex Composed of Oct-3/4 and Sox-2. Molecular and Cellular Biology 19:5453–5465. doi:10.1128/MCB.19.8.5453

58. Nishimura K, Fukagawa T, Takisawa H, Kakimoto T, Kanemaki M. 2009. An auxin-based degron system for the rapid depletion of proteins in nonplant cells. Nat Methods 6:917–922. doi:10.1038/nmeth.1401

59. Niwa H, Miyazaki J, Smith AG. 2000. Quantitative expression of Oct-3/4 defines differentiation, dedifferentiation or self-renewal of ES cells. Nat Genet 24:372–376. doi:10.1038/74199

60. Nora EP, Goloborodko A, Valton A-L, Gibcus JH, Uebersohn A, Abdennur N, Dekker J, Mirny LA, Bruneau BG. 2017. Targeted Degradation of CTCF Decouples Local Insulation of Chromosome Domains from Genomic Compartmentalization. Cell 169:930–944.e22. doi:10.1016/j.cell.2017.05.004

61. Oomen ME, Hansen AS, Liu Y, Darzacq X, Dekker J. 2019. CTCF sites display cell cycle– dependent dynamics in factor binding and nucleosome positioning. Genome Res 29:236– 249. doi:10.1101/gr.241547.118

62. Pastor WA, Liu W, Chen D, Ho J, Kim R, Hunt TJ, Lukianchikov A, Liu X, Polo JM, Jacobsen SE, Clark AT. 2018. TFAP2C regulates transcription in human naive pluripotency by opening enhancers. Nat Cell Biol 20:553–564. doi:10.1038/s41556-018-0089-0

63. Pauklin S, Vallier L. 2013. The cell-cycle state of stem cells determines cell fate propensity. Cell 155:135–47. doi:10.1016/j.cell.2013.08.031

64. Pintacuda G, Wei G, Roustan C, Kirmizitas BA, Solcan N, Cerase A, Castello A, Mohammed S, Moindrot B, Nesterova TB, Brockdorff N. 2017. hnRNPK Recruits PCGF3/5-PRC1 to the Xist RNA B-Repeat to Establish Polycomb-Mediated Chromosomal Silencing. Mol Cell 68:955–969.e10. doi:10.1016/j.molcel.2017.11.013

65. Quinlan AR, Hall IM. 2010. BEDTools: a flexible suite of utilities for comparing genomic features. Bioinformatics 26:841–842. doi:10.1093/bioinformatics/btq033

66. Raccaud M, Friman ET, Alber AB, Agarwal H, Deluz C, Kuhn T, Gebhardt JCM, Suter DM. 2019. Mitotic chromosome binding predicts transcription factor properties in interphase. Nat Commun 10:487. doi:10.1038/s41467-019-08417-5

67. Radzisheuskaya A, Chia GLB, dos Santos RL, Theunissen TW, Castro LFC, Nichols J, Silva JCR. 2013. A defined Oct4 level governs cell state transitions of pluripotency entry and differentiation into all embryonic lineages. Nat Cell Biol 15:579–590. doi:10.1038/ncb2742

68. Ramírez F, Ryan DP, Grüning B, Bhardwaj V, Kilpert F, Richter AS, Heyne S, Dündar F, Manke T. 2016. deepTools2: a next generation web server for deep-sequencing data analysis. Nucleic Acids Res 44:W160–W165. doi:10.1093/nar/gkw257

69. Rickels R, Herz H-M, Sze CC, Cao K, Morgan MA, Collings CK, Gause M, Takahashi Y-H, Wang L, Rendleman EJ, Marshall SA, Krueger A, Bartom ET, Piunti A, Smith ER, Abshiru NA, Kelleher NL, Dorsett D, Shilatifard A. 2017. Histone H3K4 monomethylation catalyzed by Trr and mammalian COMPASS-like proteins at enhancers is dispensable for development and viability. Nat Genet 49:1647–1653. doi:10.1038/ng.3965

70. Ritchie ME, Phipson B, Wu D, Hu Y, Law CW, Shi W, Smyth GK. 2015. limma powers differential expression analyses for RNA-sequencing and microarray studies. Nucleic Acids Res 43:e47. doi:10.1093/nar/gkv007

71. Robinson MD, McCarthy DJ, Smyth GK. 2010. edgeR: a Bioconductor package for differential expression analysis of digital gene expression data. Bioinformatics 26:139–140. doi:10.1093/bioinformatics/btp616

72. Sabari BR, Dall’Agnese A, Boija A, Klein IA, Coffey EL, Shrinivas K, Abraham BJ, Hannett NM, Zamudio AV, Manteiga JC, Li CH, Guo YE, Day DS, Schuijers J, Vasile E, Malik S, Hnisz D, Lee TI, Cisse II, Roeder RG, Sharp PA, Chakraborty AK, Young RA. 2018. Coactivator condensation at super-enhancers links phase separation and gene control. Science 361. doi:10.1126/science.aar3958

73. Schaffner W. 2015. Enhancers, enhancers-from their discovery to today’s universe of transcription enhancers. Biol Chem 396:311–327. doi:10.1515/hsz-2014-0303

74. Schindelin J, Arganda-Carreras I, Frise E, Kaynig V, Longair M, Pietzsch T, Preibisch S, Rueden C, Saalfeld S, Schmid B, Tinevez J-Y, White DJ, Hartenstein V, Eliceiri K, Tomancak P, Cardona A. 2012. Fiji: an open-source platform for biological-image analysis. Nat Methods 9:676–682. doi:10.1038/nmeth.2019

75. Soufi A, Donahue G, Zaret KS. 2012. Facilitators and impediments of the pluripotency reprogramming factors’ initial engagement with the genome. Cell 151:994–1004. doi:10.1016/j.cell.2012.09.045

76. Soufi A, Garcia MF, Jaroszewicz A, Osman N, Pellegrini M, Zaret KS. 2015. Pioneer transcription factors target partial DNA motifs on nucleosomes to initiate reprogramming. Cell 161:555–568. doi:10.1016/j.cell.2015.03.017

77. Stewart-Morgan KR, Reverón-Gómez N, Groth A. 2019. Transcription Restart Establishes Chromatin Accessibility after DNA Replication. Molecular Cell. doi:10.1016/j.molcel.2019.04.033

78. Strebinger D, Friman ET, Deluz C, Govindan S, Alber AB, Suter DM. 2019. Endogenous fluctuations of OCT4 and SOX2 bias pluripotent cell fate decisions. doi:10.1101/299073

79. Suter DM, Cartier L, Bettiol E, Tirefort D, Jaconi ME, Dubois-Dauphin M, Krause KH. 2006. Rapid generation of stable transgenic embryonic stem cell lines using modular lentivectors. Stem Cells 24:615–23. doi:10.1634/stemcells.2005-0226

80. Swinstead EE, Paakinaho V, Presman DM, Hager GL. 2016. Pioneer factors and ATP-dependent chromatin remodeling factors interact dynamically: A new perspective: Multiple transcription factors can effect chromatin pioneer functions through dynamic interactions with ATP-dependent chromatin remodeling factors. Bioessays 38:1150–1157. doi:10.1002/bies.201600137

81. Takaku M, Grimm SA, Shimbo T, Perera L, Menafra R, Stunnenberg HG, Archer TK, Machida S, Kurumizaka H, Wade PA. 2016. GATA3-dependent cellular reprogramming requires activation-domain dependent recruitment of a chromatin remodeler. Genome Biol 17:36. doi:10.1186/s13059-016-0897-0

82. Teves SS, An L, Hansen AS, Xie L, Darzacq X, Tjian R. 2016. A dynamic mode of mitotic bookmarking by transcription factors. Elife 5. doi:10.7554/eLife.22280

83. Wickham H. 2009. ggplot2: Elegant Graphics for Data Analysis, Use R! New York: Springer-Verlag.

84. Xiong J, Zhang Z, Chen J, Huang H, Xu Y, Ding X, Zheng Y, Nishinakamura R, Xu G-L, Wang H, Chen S, Gao S, Zhu B. 2016. Cooperative Action between SALL4A and TET Proteins in Stepwise Oxidation of 5-Methylcytosine. Mol Cell 64:913–925. doi:10.1016/j.molcel.2016.10.013

85. Xu J, Li J, Loh Y-HE, Zhang T, Jiang H, Fritzsch B, Ramakrishnan A, Shen L, Xu P-X. 2018. Brg1 controls neurosensory cell fate commitment and differentiation in the mammalian inner ear. bioRxiv 434159. doi:10.1101/434159

86. Yang Y-G, Cortes U, Patnaik S, Jasin M, Wang Z-Q. 2004. Ablation of PARP-1 does not interfere with the repair of DNA double-strand breaks, but compromises the reactivation of stalled replication forks. Oncogene 23:3872. doi:10.1038/sj.onc.1207491

87. Yuan H, Corbi N, Basilico C, Dailey L. 1995. Developmental-specific activity of the FGF-4 enhancer requires the synergistic action of Sox2 and Oct-3. Genes Dev 9:2635–2645. doi:10.1101/gad.9.21.2635

88. Zaret KS, Carroll JS. 2011. Pioneer transcription factors: establishing competence for gene expression. Genes Dev 25:2227–2241. doi:10.1101/gad.176826.111

89. Zhang Y, Liu T, Meyer CA, Eeckhoute J, Johnson DS, Bernstein BE, Nusbaum C, Myers RM, Brown M, Li W, Liu XS. 2008. Model-based analysis of ChIP-Seq (MACS). Genome Biol 9:R137. doi:10.1186/gb-2008-9-9-r137

